# A Dominant-Negative HIF-1 Isoform Protects Neuronal Development by Buffering the Response to Hypoxia in *C. elegans*

**DOI:** 10.64898/2026.08.17.745244

**Authors:** A.S. Aghabozorgi, C. Torres, K. Locsin, C. E. Carvalho

## Abstract

Animal cells respond to hypoxic stress through cell-autonomous stabilization of HIF-1, a conserved transcription factor that mediates adaptation to low-oxygen environments. Although HIF-1 is essential for viability under hypoxia, its persistent activation during development causes miswiring defects in the nervous system, raising the question of how developing neurons balance the pro-survival benefits of HIF-1 stabilization against these associated risks. Here we show that *C. elegans* neurons address this challenge by mobilizing an internal promoter within the *hif-1* locus to generate a dominant-negative isoform, HIF-1c, which acts in the nucleus to limit HIF-1/AHA-1 heterodimer formation. This buffering mechanism supports proper neuronal migration, axon guidance, and circuit formation and its loss phenocopies the defects seen when the ubiquitous HIF-1 degradation pathway is disrupted, confirming that HIF-1c serves as a critical layer of protection for the developing nervous system against hypoxic-induced errors. Lastly, we identify the retinoblastoma protein LIN-35 as a modulator of HIF-1 signaling in worms via control of *hif-1c* expression.

**Graphical Abstract:** 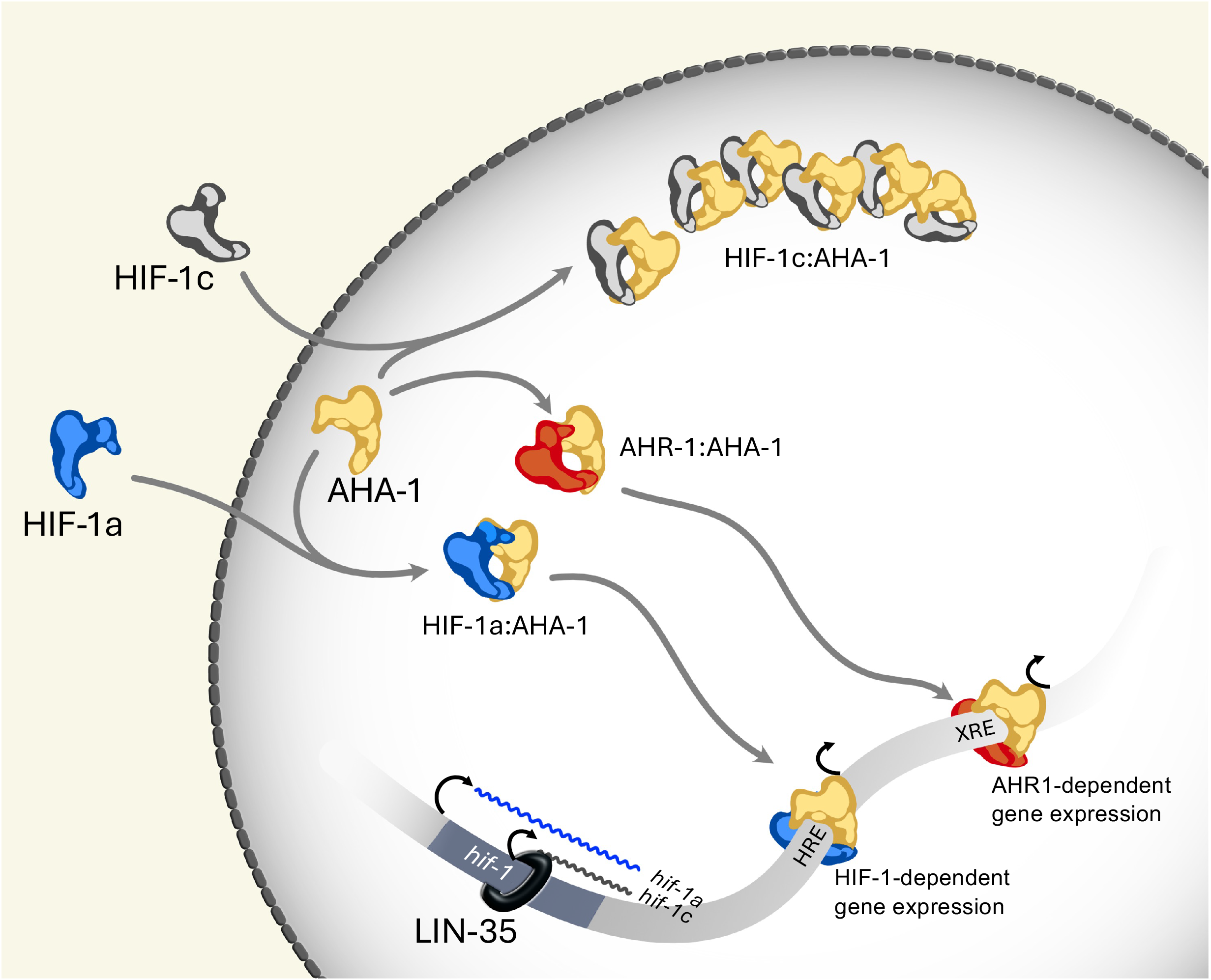

## Introduction

Metazoan development takes place under variable and often limited oxygen supply. Neurons are particularly vulnerable to hypoxia in part due to their heavy reliance on oxidative respiration and limited metabolic flexibility^1,2^. To cope with this ancient and pervasive stressor, cells deploy transcriptional programs that sense and dynamically respond to changing oxygen levels, centered on the conserved Hypoxia-Inducible Factor (HIF), a heterodimeric bHLH-PAS transcription factor composed of an oxygen-regulated α subunit (HIF-1/HIF-1α) and a constitutively expressed β subunit (AHA-1/ARNT)^3,4^. Under normoxia, HIF-1α is hydroxylated by EGL-9/PHD prolyl hydroxylases and targeted for VHL-1-dependent proteasomal degradation; under hypoxia, this degradation is blocked, allowing HIF-1α to accumulate, translocate to the nucleus, and dimerize with AHA-1 to activate genes supporting survival under low oxygen. Beyond this EGL-9/VHL-1 axis, mammals employ additional layers of HIF-1 inhibition, including tissue-specific dominant-negative HIF proteins, underscoring the need to further restrain HIF-1 signaling in specific cellular contexts^5–7^.

Despite its protective role, sustained HIF-1 activity has emerged as a direct threat to nervous system development. In a seminal study, Pocock and Hobert (2008) showed that developmental hypoxia, or genetic stabilization of HIF-1, disrupts axon pathfinding and neuronal migration in specific *C. elegans* circuits, mediated in part by transcriptional upregulation of the worm Ephrin receptor VAB-1^8^. This finding gained evolutionary significance when a similar mechanism was confirmed in zebrafish, where HIF-1-dependent upregulation of the fish Eph receptor ephrinB2a under hypoxia causes commissural axon misrouting in the forebrain^9^. Together, these studies reveal an unanticipated connection between HIF-1 activation, Eph signaling disruption and neurodevelopmental defects across vertebrates and invertebrates, suggesting a possible link between elevated HIF-1 signaling and the well-documented disruptions in axon pathfinding, synapse formation, and long-term connectivity following hypoxic injury in the developing mammalian CNS^10^.

The current understanding of HIF-1 regulation in the *C. elegans* nervous system thus presents a paradox. Like all cells, neurons rely on HIF-1 to adapt to hypoxic conditions that permeate embryonic development, yet this same transcriptional pathway appears capable of derailing the specific wiring decisions that establish neural circuits later on. How, then, do neurons maintain HIF-1 levels sufficient for survival while constraining its disruptive potential for nervous system architecture?

Here, we identify HIF-1c, a previously uncharacterized short isoform of *C. elegans* HIF-1 highly expressed in neurons via a cryptic promoter within intron 4 of the *hif-1* locus. HIF-1c retains the PAS-B dimerization domain but lacks the N-terminal bHLH DNA-binding and PAS-A domains, suggesting a protein without DNA transactivating activity but potentially able to form transcriptionally inactive complexes with AHA-1. We show that *hif-1c* expression is first detected in developing neuroblasts in parallel with transcription of the canonical *hif-1* mRNA, *hif-1a*. HIF-1c downregulates the activity of stabilized HIF-1 in neurons and hypodermal cells and loss of HIF-1c alone can phenocopy the neurodevelopmental defects observed in hypoxia and sustained HIF-1 stabilization genetic backgrounds, including HSN migration failure and PVQ axon pathfinding errors. We extend the list of HIF-1-sensitive neurons in *C. elegans* by describing defects in the connections between VC4/VC5 postembryonic neurons, part of the egg-laying circuit. Connectivity defects in VC4/VC5 are observed in *vhl-1* and *hif-1c* mutants and as other HIF-1-driven neuronal defects, are mediated by Eph signaling. Mechanistically, we propose a model in which HIF-1c antagonizes HIF-1 activity by sequestering AHA-1 in the nucleus, reducing the pool of AHA-1 available to form productive HIF-1a/AHA-1 complexes. Finally, we implicate the pRB homolog and neuronal fate regulator, LIN-35, as an upstream regulator of *hif-1c* expression in neurons whose activity is required to prevent HIF-1-driven neurodevelopmental defects. These findings extend the regulatory network controlling HIF-1 activity in *C. elegans* and implicate HIF-1c as a novel tissue-specific inhibitor responsible for keeping HIF-1 signaling within physiological limits during developmental hypoxia, thereby ensuring correct wiring of the nervous system.

## Results

### An internal promoter within the *hif-1* locus mediates expression of a neuronally enriched HIF-1 isoform

To investigate potential differences in HIF-1 signaling in neurons, we focused on the transcriptional products of the sole *hif-1* gene in *C. elegans* (F38A6.3). *hif-1* is predicted to generate at least six mRNAs. The ubiquitously expressed HIF-1a isoform encoded by *hif-1a* mRNA corresponds to the canonical bHLH-PAS transcription factor in worms and, like its vertebrate HIF-1α homologs, is required for adaptation to hypoxia^11^. Four of the remaining annotated *hif-1* mRNAs (*hif-1b,d,e* and *f*) are splice variants encoding isoforms predicted to retain the N-terminal bHLH domain (Figure 1A). The exception is *hif-1c*, a transcript encoding a HIF-1 isoform lacking the DNA binding domain but potentially capable of forming transcriptionally inactive heterodimers with AHA-1/ARNT via the remaining PAS-B domain. The annotated *hif-1c* mRNA contains a unique start codon in a cryptic exon immersed within intron 4 about 100bp downstream of a mapped transcription start site (TSS), suggesting it may originate through an independent transcriptional event^12,13^. We therefore determined whether intron 4 harbors an internal promoter responsible for *hif-1c* expression using a reporter approach. Worms stably co-expressing transcriptional reporters driven by the canonical promoter (*Phif-1::nRFP* as a proxy for *hif-1a* expression) or intron 4 sequences (*Pi4::GFP* as a proxy for *hif-1c* expression) were produced. In striking contrast to the ubiquitous expression of *Phif-1::nRFP*, strong GFP signal is observed primarily across the *C. elegans* nervous system where it marked all major ganglia as well as nerve cords, lateral processes and commissures (Figure 1B; Figure S1B). Importantly, neurons expressing *Pi4::GFP* also express *Phif-1::nRFP* (Figure 1C). Developmentally, signals from both reporters are first observed in neuroblasts during ventral enclosure at a time of intense cell migration when the basic architecture of the worm nervous system is being laid out. As embryonic development progresses, *Phif-1::nRFP* expression extends to other cells of the embryo, whereas *Pi4::GFP* signal becomes enriched in differentiated neurons (Figure S2). In addition to neurons, *Pi4::GFP* signal is detected in the distal tip cell (DTC) of the somatic gonad and uv1 neuroendocrine cells. Weak expression is also observed in the hypodermis, muscle and intestine (Figure S1A). Notably, not all neurons appear to utilize the internal promoter. Notable exceptions are neurons of the egg-laying circuit VC4, VC5 and HSN (Figure S1B,C). These results are consistent with the existence of an internal promoter in intron 4 of *hif-1* (hereafter referred to as Pi4), presumably important for tissue-specific expression of *hif-1c,* and suggest that Pi4 becomes active precisely at the onset of neuronal differentiation and circuit formation.

**FIGURE 1:**
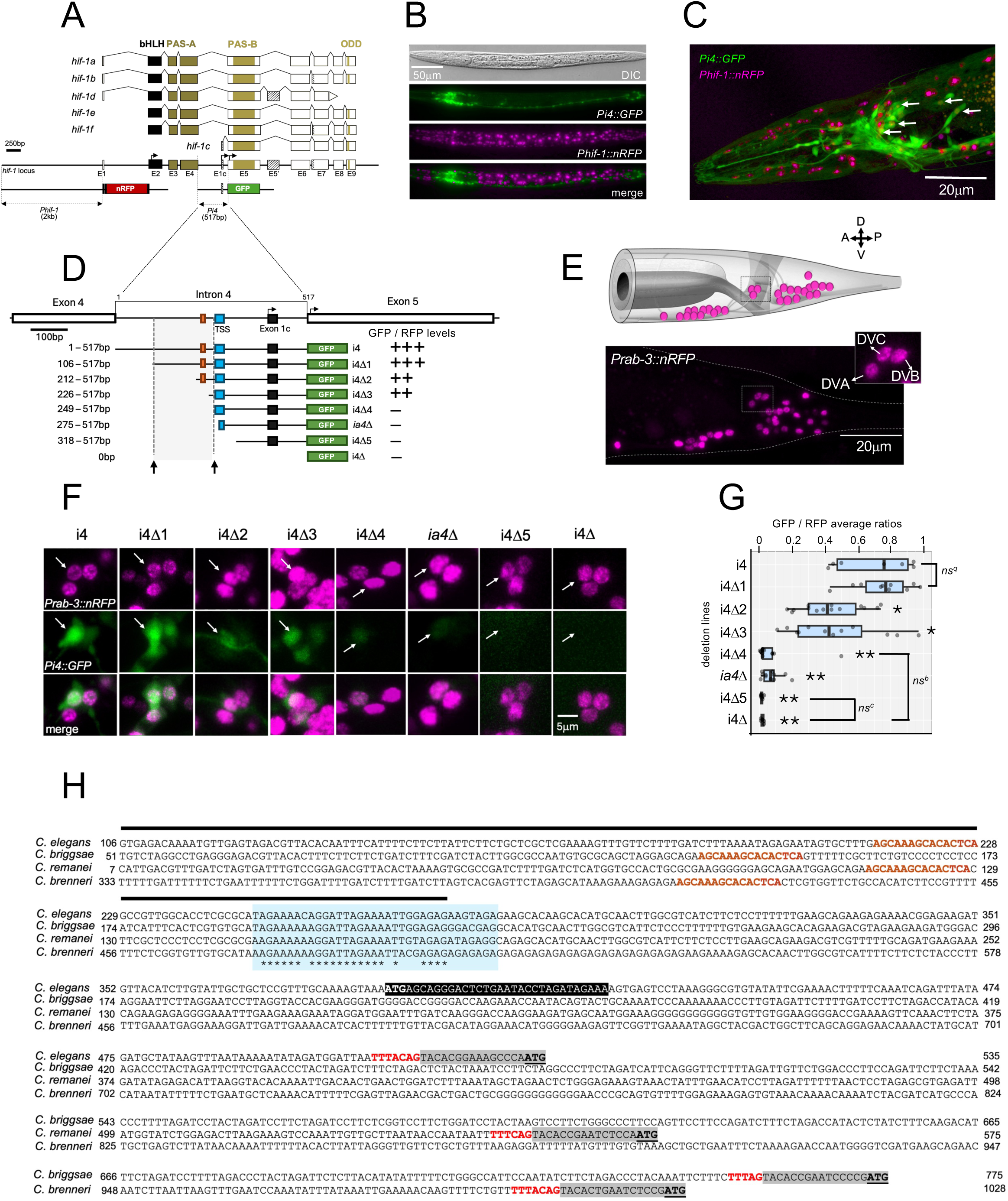
A short *hif-1* transcript (*hif-1c*) is expressed via an internal promoter within intron 4. (A) *C. elegans hif-1* locus and transcripts with protein domains as colored boxes and alternative spliced region as striped boxes; E1c denotes *hif-1c*’s cryptic exon 1. Transcriptional reporters for the canonical promoter (*Phif-1*::*nRFP*) and intron 4 (*Pi4::GFP*) are shown below. (B) L2 larva co-expressing *Phif-1*::*nRFP* and *Pi4::GFP* (*sasIs7*); anterior left. (C) Head of an adult *sasIs7* worm; arrows indicate nerve ring neuronal nuclei co-expressing the two reporters. (D–G) Deletion analysis of Pi4. (D) Intron 4 region with putative TSS as a blue box and the tested Pi4 deletion GFP constructs shown below. Relative GFP expression levels are indicated on the right. Arrows mark a 143-bp region (bp 106-249) sufficient for *Pi4::GFP* expression. (E) Neuronal cell body positions in the tail (modified from WormAtlas), with the corresponding tail region of an L4 worm expressing *Prab-3*::*nRFP* below; inset shows dorsorectal ganglion neurons (DVA, DVB, DVC) used for quantification. (F) Dorsorectal ganglia in transgenic worms expressing Pi4 deletion GFP reporters; arrows indicate DVC neurons quantified. (G) GFP/RFP ratios in DVC neurons, averaged across two independent lines per construct. *p<0.05, **p<0.001, nsᵃ p=0.38 vs. positive control (i4); nsᵇ p=0.24, nsᶜ p=0.29 vs. negative control (i4Δ). (H) Sequence alignment of the internal promoter across *Caenorhabditis* species. The conserved TSS (blue) is 57% identical across all four species (asterisks); the 15-bp Box1 sequence (brown font in D) is marked in brown fonts and the region deleted in the *ia04* allele is marked by a dark bar above. E1c and its putative first ATG (black highlight) in *C. elegans* intron 4 are not conserved in other species, unlike the in-frame ATG in exon 5 (grey highlight) and the SL1-like sequence (red) at the end of intron 4.

To map the boundaries of Pi4, we performed a reporter deletion analysis by sequentially deleting ∼100 bp of 5’ sequence across the entire 517 bp of intron 4. (Figure 1D). Fluorescence levels were measured from the dorsorectal ganglion neuron DVC (Figure 1E), normalized to *Prab-3::RFP* fluorescence and compared with *Pi4::GFP* control lines (construct i4). As shown in Figure 1F and quantified in Figure 1G, removing the first 106bp of intron 4 (construct i4Δ1) did not affect expression levels (p=0.38). In contrast, progressively deleting sequences immediately upstream of the putative TSS reduced GFP expression significantly (construct i4Δ2; p=0.01 and construct i4Δ3; p=0.04). Transgenic worms carrying reporters that retained exclusively the TSS (construct i4Δ4) as well as downstream regions (construct i4Δ5), including the putative exon 1 of *hif-1c* (E1c), showed no significant reporter transactivation as compared to the control promoterless GFP reporter lines (i4Δ; p=0.24 and p=0.29, respectively). Moreover, a reporter that replicates the 3’ deletion boundary of the reference *hif-1* allele *ia4* (*ia4*Δ) by removing the first 275bp of intron 4, including part of the TSS, showed a significant reduction in GFP expression as compared to the positive control (p=0.0005). To identify possible functionally important sequences in Pi4, we assessed sequence conservation in intron 4 of other *Caenorhabditis* species (Figure 1H). As expected, the 35bp TSS region is highly conserved (63% identity) among *C. elegans, C. briggsae, C. remanei and C. brenneri*. A 100% identical 15bp sequence within Pi4 (termed box1) is also present across the three non-*elegans* species, though its removal in construct i4Δ3 does not significantly affect expression in *C. elegans* (p=0.69 as compared to i4Δ2; Figure 1D,G and H). Finally, the putative E1c sequence of *C. elegans hif-1c*, including its first ATG, is not conserved in intron 4 of the other three *Caenorhabditis* species. Taken together, these results narrowed down Pi4 to a 143bp critical region upstream of the conserved TSS in intron 4 (positions 106 to 249) that is required and necessary for expression.

### HIF-1c is a transcriptionally inactive and oxygen-sensitive

Cells tightly control nuclear HIF-1 activity through a conserved oxygen-dependent degradation machinery^14^. We generated a translational reporter to investigate the oxygen sensitivity and subcellular localization of the HIF-1 isoform expressed via Pi4. The genomic region from intron 4 to exon 9 was fused in frame to GFP and used to generate *Pi4::hif-1c::GFP* expressing worms (*sasIs6;* Figure 1A). In these worms, GFP signal is concentrated in the nuclei of neurons, though a weak cytoplasmic signal in cell bodies and dendrites is sometime present (Figure 2B,C; Video S1). Consistent with the presence of an ODD domain, the signal increases in wild-type worms incubated in hypoxia (1% O_2_; Figure S3, Video S2) and in mutants with compromised oxygen-dependent degradation of HIF-1 (Figure 2D,E; p=0.004). To verify the exact transcript(s) expressed by Pi4, *sasIs6*-specific cDNA was amplified using a GFP-specific 3’ primer and an SL1 splice leader primer that tags the 5’of most mRNAs in *C. elegans* (Figure 2A). Cloning and sequencing confirmed that *sasIs6* expresses exclusively *hif-1c* mRNAs lacking the predicted E1c sequence and the E1c ATG, removed during SL1 transplicing (Figure S4A,B). The removal of the cryptic exon sequences from the processed *hif-1c* mRNA is consistent with the lack of conservation of E1c in intron 4 of other species (Figure 1H). Instead, HIF-1c initiates translation from Met233 of HIF-1a, an in-frame methionine encoded by exon 5 (Figure S4C). Collectively, these results confirmed that neurons utilize Pi4 to produce HIF-1c, a shorter, oxygen-regulated HIF-1 isoform that lacks the N-terminal bLHL and PAS-A domains.

**FIGURE 2:**
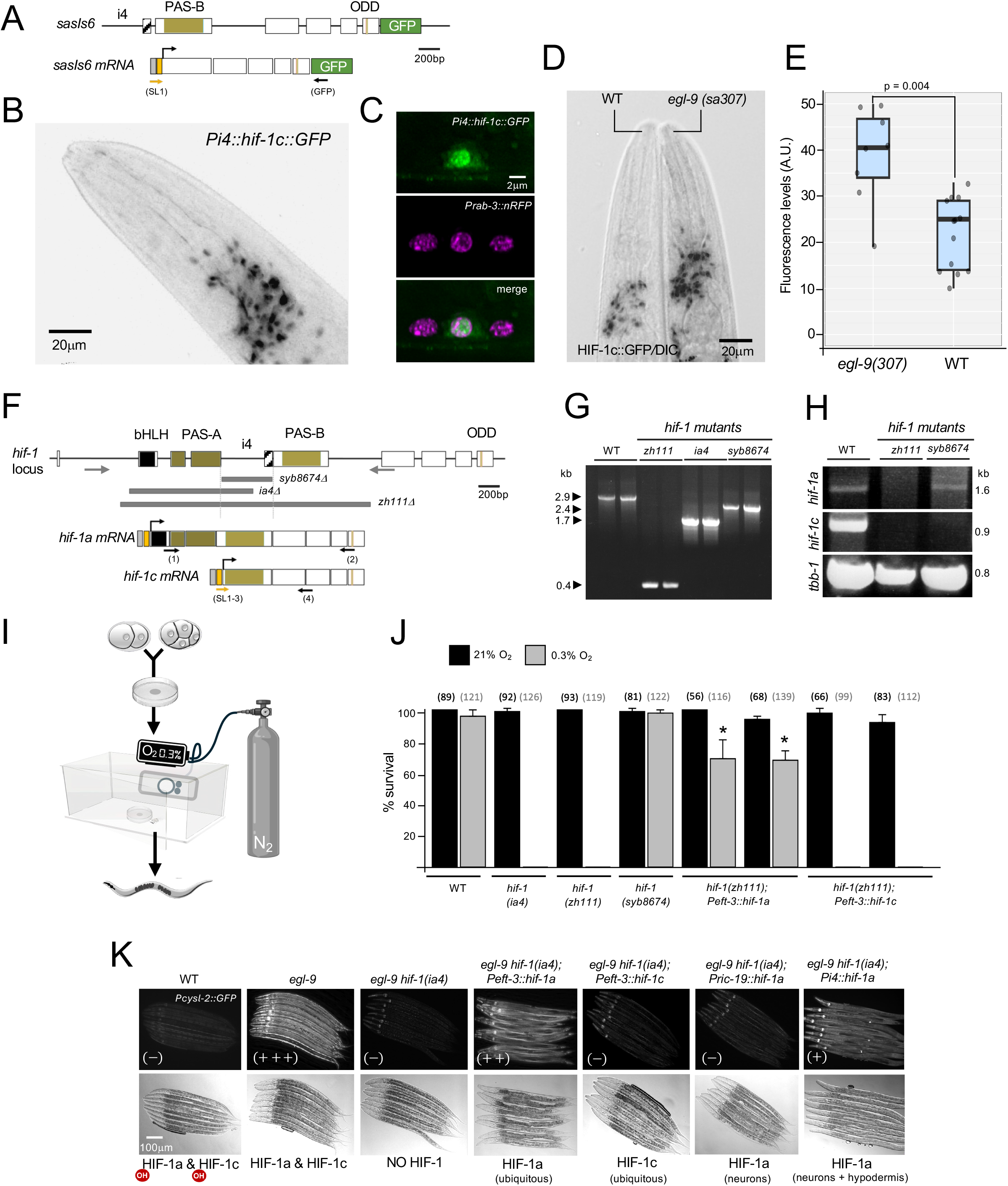
HIF-1c is an oxygen-sensitive HIF-1 isoform not required for adaptation to hypoxia. (A) *Pi4::hif-1c::GFP* translational reporter (*sasIs6*); cDNA primers used to specifically isolate *hif-1c*::GFP transcripts are shown below. (B) DIC/inverted fluorescence image of an *egl-9(sa307)* adult head expressing HIF-1c::GFP, visible in neuronal cell bodies of the anterior ganglia and nerve ring. (C) VNC motor neuron cell body expressing HIF-1c::GFP. (D) DIC/HIF-1c::GFP overlay of the head region of *egl-9(sa307)* and WT worms. (E) Quantification of HIF-1c::GFP in the nerve ring of WT and *egl-9(sa307)* worms (n=26). (F) Deletion alleles used in this study: *zh111* (2,464 bp), *ia04* (1,200 bp), and *syb8674* (517 bp), with genotyping (grey arrows) and transcript-specific (black arrows) primers shown. Top arrows mark start codons for *hif-1a* and *hif-1c*; yellow boxes indicate the transpliced SL1 site in *hif-1a* and *hif-1c* mRNAs; grey boxes indicate 5’UTRs. (G) Representative single-worm genotyping for WT (N2) and the three *hif-1* deletion mutants. (H) cDNA amplification of *hif-1a*- and *hif-1c*-specific amplicons using primers from (F), with *tbb-1* cDNA as control. (I) Experimental layout for the hypoxia sensitivity assay. (J) Embryonic viability of WT, *hif-1*, and rescue lines expressing HIF-1a or HIF-1c transgenes; total embryos scored (n) across three independent experiments shown above each bar. p<0.001 relative to zh111 at 1% O₂. (K) *Pcysl-2*::*GFP* (*nIs470*) expression across genetic backgrounds with variable HIF-1 stabilization; predicted HIF-1 isoform produced in each background shown below. OH indicates hydroxylation.

The intron 4 of *hif-1* is removed by *zh111*, a large deletion allele that, as *ia4*, impacts other *hif-1* mRNAs in addition to *hif-1c* ^15^ (Figure 2F). To dissect the specific function of HIF-1c in worms, we sourced an allele that disrupts HIF-1c exclusively. *hif-1(syb8674)* is a CRISPR/Cas9-generated deletion that removes all 517 bp of intron 4 which eliminates Pi4 and fuses exons 4 and 5 in-frame (Figure 2F,G; Figure S4D). *hif-1c* transcription is completely abrogated in *hif-1(syb8674)* worms, confirming that HIF-1c production entirely relies on Pi4. Importantly, *hif-1a* expression is unchanged in these animals, indicating that *hif-1(syb8674)* worms retain intact HIF-1-dependent signaling (Figure 2H). The expression of an additional HIF-1 isoform in neurons could reflect a heightened demand for HIF-1 in a tissue highly sensitive to hypoxia. However, the absence of a DNA-binding domain in HIF-1c argues against a transcriptional role of this isoform. To test whether HIF-1c nonetheless performs a HIF-1-like function, we examined the viability of *hif-1(syb8674)* embryos in hypoxia assays (Figure 2I). As with *hif-1(zh111)* and *hif-1(ia04)* worms, which lack HIF-1 activity*, hif-1(syb8674)* animals are viable and display no overt phenotypes in room air (p=0.42 as compared to wild type). Unlike *hif-1(zh111)* and *hif-1(ia04)*, however, *hif-1(syb8674)* embryos also develop normally at 0.3% oxygen, suggesting that HIF-1c is dispensable for hypoxic adaptation and confirming that these animals can mount HIF-1-dependent responses (p=0.38 as compared to wild type). Accordingly, while ubiquitous expression of canonical HIF-1a using the *eft-3* promoter rescues the hypoxia-induced lethality of *hif-1(zh111)* mutants (p<0.001), ectopically expressing HIF-1c in all cells of *hif-1(zh111)* embryos failed to reduced lethality under low-oxygen conditions (Figure 2J). HIF-1c may contribute only minimally to HIF-1 activity below the sensitivity of the embryonic viability assay. We corroborated these results by assessing the impact of HIF-1c in the expression of *Pcysl-2::GFP*, a bona fide *hif-1* reporter carrying the promoter of the cysteine synthase 2 gene^16^. As shown in Figure 2K, *Pcysl-2::GFP* expression increases several-fold in the hypodermis of *egl-9* mutants compared to wild type, consistent with HIF-1 accumulation in this background. Induction of the reporter is entirely dependent on HIF-1 activity, as it is suppressed in *egl-9 hif-1(ia04)* double mutants. In these worms, *Peft-3*-driven expression of HIF-1a, but not HIF-1c, restores reporter induction, confirming that HIF-1c lacks transactivation activity.

Ectopic expression of HIF-1a pan-neuronally via the *ric-19* promoter fails to increase *Pcysl-2::GFP* fluorescence in the hypodermis, reflecting a cell-autonomous requirement for HIF-1 in this tissue. Indeed, driving HIF-1a using the internal promoter (*Pi4::hif-1a*), which is active in both neurons and hypodermal cells (Figure S1A), upregulates hypodermal *Pcysl-2::GFP* expression, validating the results with *Peft-3* transgenes. These findings argue against a transcriptional role for HIF-1c and suggest it does not positively contribute to HIF-1 signaling in *C. elegans*

### HIF-1c antagonizes HIF-1 signaling

Having established that HIF-1c does not act as a HIF transcriptional activator, we next asked whether it might instead downregulate HIF-1 signaling. We reasoned that tissue-specific HIF-1c may be necessary to restrain HIF-1 levels by interfering with HIF-1/AHA-1 heterodimer assembly or function. If so, loss of HIF-1c in *hif-1(syb8674)* animals would be expected to produce measurable increases in HIF-1 activity. Indeed, *Pcysl-2::GFP* expression is significantly upregulated in the hypodermis of *hif-1(syb8674)* worms, despite an intact oxygen-dependent HIF-1 degradation pathway in these mutants (p<0.05; Figure 3A). This increase is more modest than that observed in *egl-9(sa307)*, confirming that in normoxia, HIF-1 signaling is predominantly kept in check through proteasome activity (Figure 3A,B). The increase in HIF-1 activity observed in the hypodermis of *hif-1c* mutants should be mirrored by a corresponding reduction in HIF-1-mediated transcription in animals overexpressing HIF-1c. As expected, *Pcysl-2::GFP* induction is attenuated in the hypodermis of *egl-9* mutants over expressing HIF-1c, suggesting that HIF-1c acts to counteract the activity of stabilized HIF-1 (p<0.001; Figure 3C,D). The buffering effect of HIF-1c on HIF-1 signaling is not a reporter artifact, since it can be replicated using *Pnhr-57::GFP*, an independent HIF-1-regulated transgene (Figure S5). Given that *hif-1c* is predominantly expressed in the nervous system, we sought to determine whether its antagonistic role in regulating HIF-1 activity extends to neurons. HIF-1 stabilization has previously been shown to upregulate tryptophan hydroxylase (*tph-1*) in ASG sensory neurons, driving increased serotonin production to activate a cryptic neuronal circuit that modulates chemotaxis behaviour in hypoxic environments^17^. Consistent with this, constitutive HIF-1 activation in *vhl-1(ok161)* mutants results in induction of a *tph-1* transcriptional reporter in ASG neurons. Elevated *Ptph-1*::GFP expression is also observed in ASG of *hif-1(syb8674)* animals under normoxic conditions, reflecting increased HIF-1 activity in these neurons (Figure 3E,F). Since ASG neurons express *hif-1c* (Figure 3G), loss of HIF-1c activity in these cells likely underlies the observed increase in HIF-1-dependent gene expression. Finally, we examined whether HIF-1c influences the egg-laying defect (Egl) of *egl-9* mutants, a well-established proxy phenotype for HIF-1 activity in worms. Elevated HIF-1 levels in neurons and uv1 cells, two cell types with high *hif-1c* expression (Figures S1A,B), underlies the egg-retention behaviour of *egl-9* worms and has been previously exploited to identify downstream regulators of HIF-1 in *C. elegans*^18–20^. Abolishing HIF-1 activity in *egl-9(sa307) hif-1(ia4)* double mutants fully suppresses the Egl defect (p<0.001; Figure 3H). In contrast, the *hif-1(syb8674)* allele, which depletes HIF-1c specifically, fails to restore normal egg-laying in *egl-9* mutants (p=0.621), confirming that HIF-1c stabilization does not contribute to increased HIF-1 activity in these worms. Conversely, overexpression of HIF-1c in *egl-9; sasIs6* animals significantly increases egg-laying despite impaired HIF-1 degradation (p<0.001; Figure 3H,I). Collectively, these results establish an inhibitory role for HIF-1c in modulating HIF-1 activity across both neuronal and non-neuronal cell types.

**FIGURE 3:**
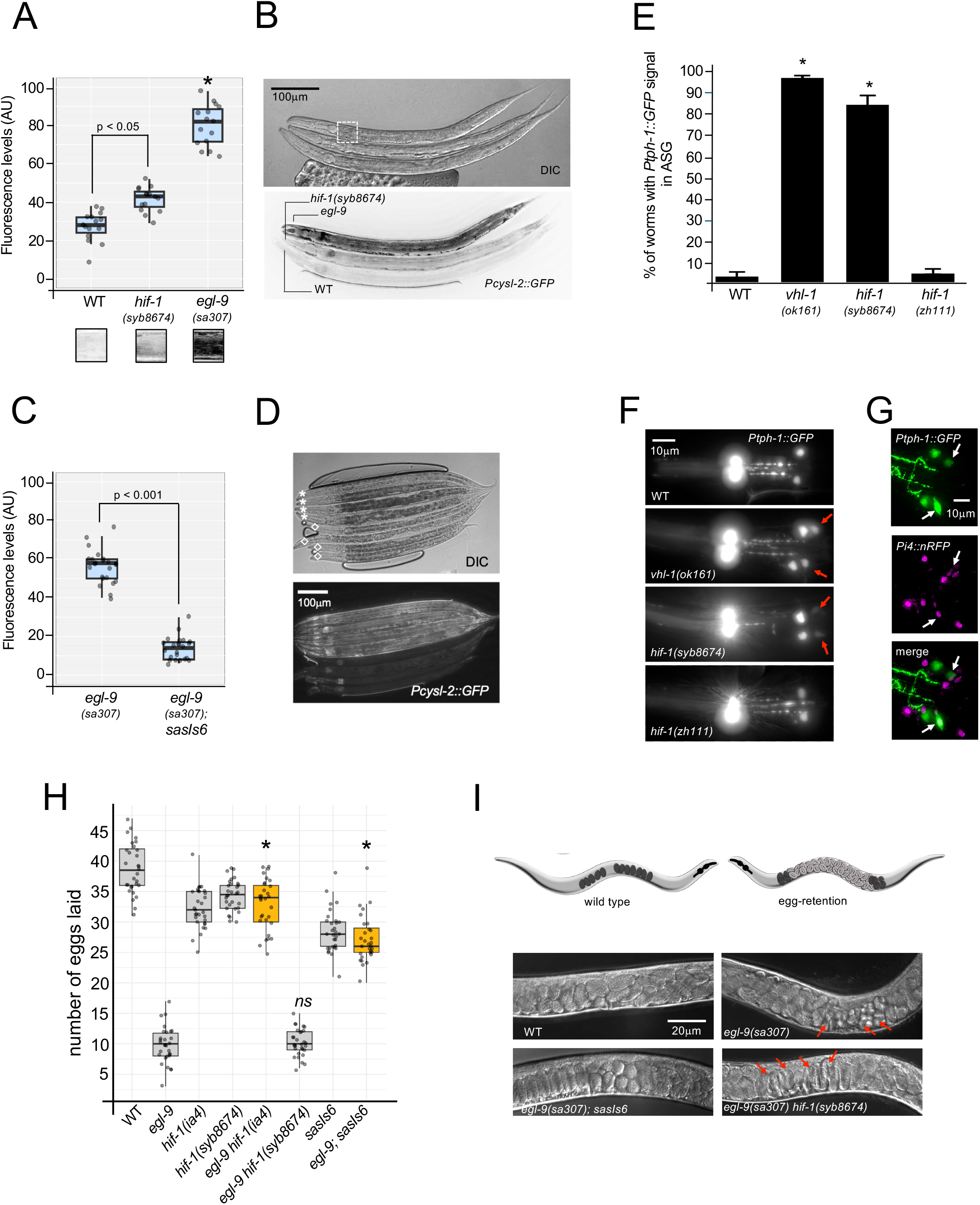
HIF-1c antagonizes HIF-1-dependent gene expression. (A,B) Loss of HIF-1c increases HIF-1-dependent gene expression. (A) Quantification of *Pcysl-2*::*GFP* fluorescence in the hyp7 syncytium of WT (*nIs470*), *egl-9(sa307)*, and *hif-1(syb8674)* worms; representative images in (B), with region used for fluorescence quantification boxed and replicated below the graph. (C,D) HIF-1c activity reduces HIF-1-dependent expression in the hypodermis. (C) Quantification of *Pcysl-2*::*GFP* signal changes in *egl-9* mutants versus *egl-9; sasIs6* worms; representative images in (D), with *egl-9(sa307)* marked by asterisks and *egl-9(sa307); sasIs6* by diamonds. (E,F) *Ptph-1*::*GFP* (*zdIs13*) expression in ASE neurons of worms with increased HIF-1 signaling; quantification in (E) and representative images in (F), arrows indicating the two ASE neurons. p<0.001 vs. *zdIs13* control. (G) Co-localization of *Pi4::nRFP and Ptph-1::GFP* in ASE (arrows). (H) HIF-1c overexpression suppresses HIF-1-dependent egg-laying defects in *egl-9* mutants. Eliminating HIF-1 activity (*ia04*) or increasing HIF-1c activity (*sasIs6*) significantly suppresses the *egl-9* egg-laying defect (*p<0.0001, yellow boxes), whereas *syb8674* does not (ns; p=0.26). (I) Representative DIC images of the uterus in day-2 adults of the indicated genotypes; red arrowheads mark late-stage, three-fold embryos retained in the uterus.

### HIF-1c is required for HSN migration, PVQ axon pathfinding and VC4/5 connectivity

The neuronal enrichment of HIF-1c, combined with its ability to antagonize HIF-1 activity, makes it an ideal candidate for protecting neurons from neurodevelopmental defects arising from sustained HIF-1 stabilization. To test this, we investigated whether the loss of HIF-1c results in defects akin to those observed in worms with high HIF-1 activity. Consistent with this, *hif-1(syb8674)* neurons exhibit axon pathfinding and migration defects similar to those in worms developed under hypoxia, in *vhl-1* or *egl-9* mutants and in transgenic animals expressing HIF-1^P621A^, an oxygen-insensitive variant with a disrupted degradation domain^8^. Specifically, in over 25% of *hif-1c* mutants, axons of the two PVQ interneurons intermittently leave their respective fascicles in the ventral nerve cord and abnormally cross the midline (p<0.001; Figure 4A-C). Similarly, embryonic migration of HSN neurons from the tail to the vulva region, where they later innervate the egg-laying circuit, is disrupted in 18% of *hif-1(syb8674)* animals (p<0.001; Figure 4E-G). In these animals, one HSN neuron fails to complete its anterior migration, with the cell body remaining ectopically in the tail from where it extends processes toward the ventral nerve cord (VNC). Expressing a *hif-1c* rescue transgene under driven by Pi4 rescues both PVQ and HSN defects (p<0.001; Figure 4C,G). While both PVQ and HSN neurons display developmental defects consistent with loss of HIF-1c function, *hif-1c* expression is detected in PVQ but not in HSN, suggesting that HIF-1c may have both cell-autonomous and non-autonomous roles in regulating neurodevelopment (Figure 4D,H).

**FIGURE 4:**
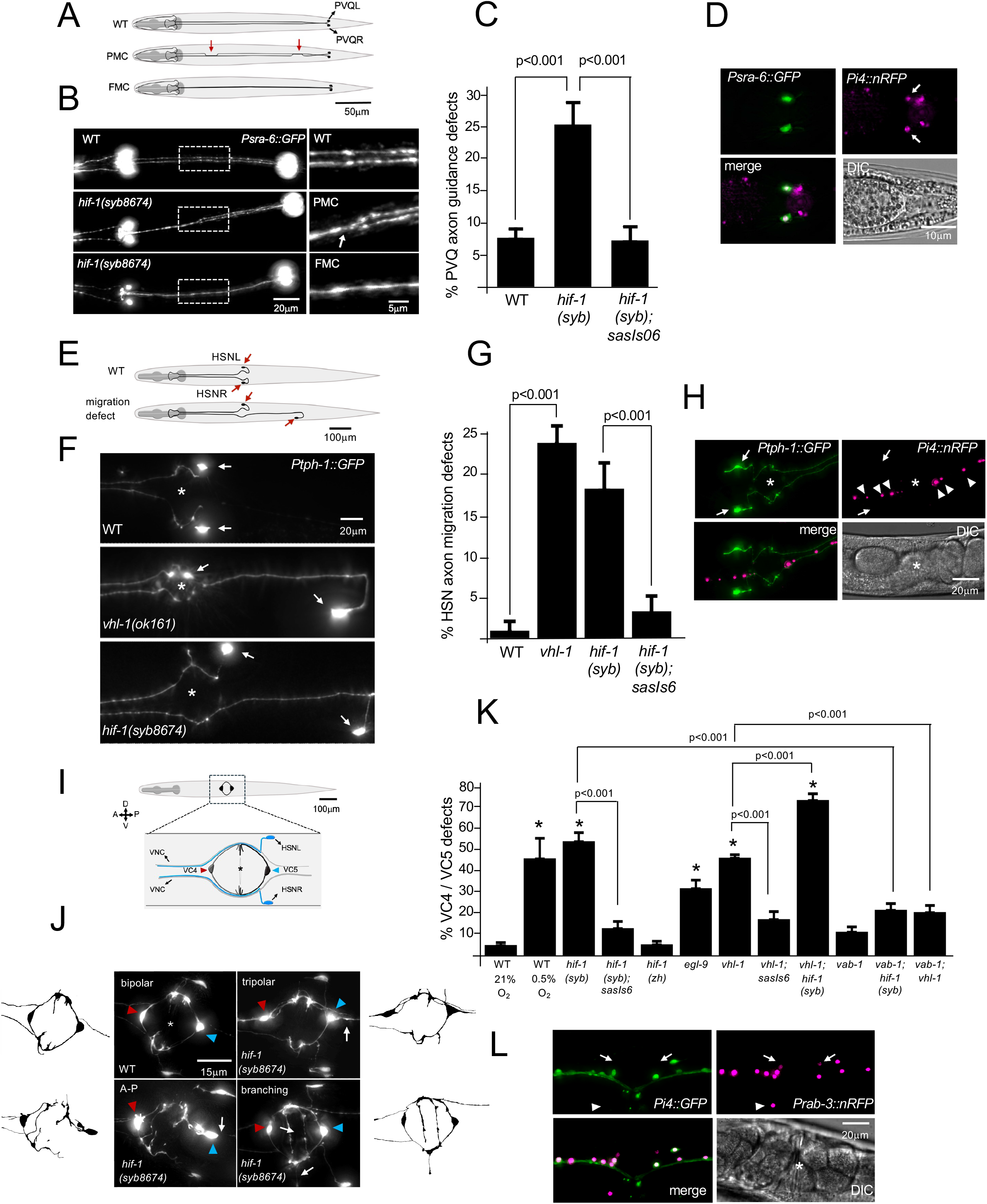
HIF-1c depletion leads to neuronal defects caused by increased HIF-1 signaling. (A-D) HIF-1c is required for PVQ axon guidance. (A) L1 larva showing PVQ axon processes along the two ipsilateral VNC tracks in WT (top) versus *hif-1(syb8674)* mutants with varying degrees of axon guidance defects: partial midline crossing (PMC) and full midline crossing (FMC); arrows mark misrouted sections. (B) L1 larvae expressing the PVQ reporter *Psra-6*::*GFP* (*oyIs14*) along the VNC; boxes and insets mark the central VNC region where two parallel axon tracks are seen in WT, with PMC and FMC defects in *hif-1(syb8674)* shown below (arrow indicates axon crossing). (C) Quantification of PVQ axon guidance defects in L1 larvae. (D) Co-localization of *Psra-6*::*GFP* and *Pi4::nRFP* in PVQ neurons (arrows). (E-H) Defects in HSN migration. (E) L4 larva showing position of the two HSN neurons (arrows). (F) HSN neurons (arrows) expressing *Ptph-1*::*GFP* (*zdIs13*), correctly migrated in WT (top) but shows defects in *vhl-1(ok161)* and *hif-1(syb8674)*, quantified in (G). (H) HSN neurons do not express *Pi4::nRFP* (arrows); instead, HIF-1c appears to be produced by ventral cord neurons in this region (arrowheads). (I-L) HIF-1c regulates proper wiring of the egg-laying circuit at the L4 stage. (I) Anatomical position of VC4/5 neurons around the vulva (asterisk). (J) Insets of VC4/5 and HSN neurons expressing P*cat-1*::GFP (*otIs221*), with trace diagrams alongside for clarity; VC4 and VC5 are indicated by red and blue arrowheads, respectively. The stereotypical bipolar arrangement of VC4/5 axons connecting laterally around the vulva (asterisk) is shown top left (WT); representative defects (extra axons/tripolar, A-P polarity, terminal branching) in *hif-1(syb8674)* VC4 and VC5 neurons are indicated (arrows). (K) Quantification of VC4/VC5 neuronal defects; averages of three experiments shown (n ≥ 60 worms/experiment); *p<0.001 vs. WT at 21% oxygen. Complete genotypes on the x-axis: *hif-1(zh111); hif-1(syb8674); egl-9(sa307); vhl-1(ok161); vab-1(ok1699)*. (L) *hif-1c* is not expressed in VC4 and VC5 neurons; VC4/5 (arrows) expressing *Prab-3::nRFP*, arrowheads mark a HSN.

PVQ axon pathfinding and HSN cell body migration occur in the context of the hypoxic embryo. Yet, *hif-1c* expression in the nervous system persists beyond embryonic development through larval stages and into adulthood, when animals navigate environmental oxygen gradients. This raises the question of whether intermittent hypoxia also threatens the post-embryonic nervous system. The egg-laying circuit represents one of the last neuronal circuits to be functionally assembled in *C. elegans*, with HSN and VC neurons undergoing final differentiation and synaptogenesis at the L4 stage. During vulval development, VC4 and VC5 extend processes dorsally around the vulval slit, producing a distinctive bipolar morphology unique among VNC motor neurons that depends on polarity cues from neighboring neurons and branching-inhibitory signals from the vulval epithelium, respectively ^21–24^. These processes meet laterally at the vulval midline and, together with HSN axonal branches, innervate the vulval muscles to complete the egg-laying circuit formation (Figure 4I,J). We asked whether elevated HIF-1 activity during late L4 disrupts formation of the egg-laying circuit. Stabilizing HIF-1 in wild-type animals exposed to hypoxia, or in *vhl-1* and *egl-9* mutants, results in a spectrum of VC4/VC5 connectivity defects (p<0.001; Figure 4K). These include the presence of supernumerary processes producing a tripolar rather than bipolar morphology, ectopic neurites that aberrantly cross the vulval midline and outgrowth defects when VC4/VC5 cell bodies fail to project dorsally and instead extend processes along the anterior/posterior axis of the VNC, resembling the morphology of the non-vulval VC neurons. Consistent with a post-embryonic role for HIF-1c in restraining HIF-1 activity, approximately 55% of *hif-1(syb8674)* animals exhibit VC4/VC5 connectivity defects, which are rescued by restoring *hif-1c* expression using its endogenous promoter (p<0.001; Figure 4J,K). These neuronal defects arise from elevated HIF-1 signaling, as they are fully suppressed in *hif-1(zh111)* null animals lacking not only HIF-1c, but all other HIF-1 isoforms (p=0.774 as compared to the normoxia control). Supporting a novel mechanism of HIF-1 inhibition in *C. elegans*, overexpression of HIF-1c in *vhl-1* mutants significantly reduces the penetrance of VC4/VC5 wiring defects (p<0.001), while combined loss of *hif-1c* in *hif-1(syb8674); vhl-1* double mutants further increases the proportion of animals affected (p<0.001; Figure 4K). Together, these genetic interactions indicate that HIF-1c modulates the activity of stabilized HIF-1, independently of the canonical HIF-1 degradation machinery.

In both *C. elegans* and zebrafish, the defects resulted from HIF-1 activation are mediated in part through Eph signaling^8,9^. *vab-1*, the sole Eph receptor in worms, is a direct target of HIF-1 and a key regulator of neuronal migration and axon pathfinding. We find that VAB-1 similarly meidates HIF-1-dependent defects in VC4/5. While eliminating Eph signaling in *vab-1* mutants causes only a subtle but significant increase in VC4/5 wiring defects (p<0.031; Figure 4K), depletion of VAB-1 in *hif-1(syb8674)* and *vhl-1* mutant backgrounds strongly suppresses the defects in VC4/5 (p<0.001; Figure 4K). In the *C. elegans* embryo, increased VAB-1 expression in ventral motor neurons (MNs) non-autonomously disrupts PVQ axon guidance along the VNC by interfering with the establishment of a molecular Ephrin barrier in the midline^25^. We found that VC4/5, like HSN, do not express *hif-1c*, consistent with local Eph dysregulation arising from exacerbated VAB-1 displayed on the surface of neighboring cells instead (Figure 4L). Collectively, these findings support a model in which HIF-1c attenuates HIF-1 activity in the ventral midline by acting on MNs to ensure proper local Eph signaling levels required for neuronal development.

### HIF-1c binds to AHA-1 in the nucleus

How does HIF-1c mechanistically downregulate HIF-1 signaling? Tissue-specific dominant-negative HIF-3α isoforms that reduce the formation of viable HIF-1 heterodimers have been previously described in mice and humans^5–7,26^. These proteins associate with HIF-1α or HIF-1β subunits, either preventing their nuclear translocation or forming transcriptionally unproductive nuclear complexes that deplete the subunits available for active HIF-1α/β heterodimer formation. In principle, since HIF-1c is localized to both the cytoplasm and nuclei of neuronal cell bodies, it could act through either mechanism. Using bimolecular fluorescence complementation (BiFC) with split-Venus constructs (VN173 and VC155), we tested whether HIF-1c physically interacts *in vivo* with HIF-1a or AHA-1 (Figure 5A). Ubiquitous expression of HIF-1a and AHA-1 constructs via the *Phsp-16* heat shock promoter produced robust nuclear fluorescence in intestinal cells of *vhl-1* and *egl-9* mutants (Figure 5B,C). Similarly, HIF-1c/AHA-1 interaction was observed exclusively in the nucleus of both intestinal cells and neurons and depended on the remaining PAS domain of HIF-1c (PAS-B) as a HIF-1c mutant lacking PAS-B failed to complement with AHA-1 (Figure 5A-C; Figure S6). Association of HIF-1a/c with AHA-1 in the nucleus appear independent of HIF-1 hydroxylation as observed by complementation in the intestine of *egl-9*, a HIF-1 hydroxylation-deficient mutant. We found no evidence of HIF-1c/HIF-1a dimerization, indicating that HIF-1c does not directly inhibit HIF-1a function. Importantly, Venus complementation involving HIF-1c constructs was never detected in the cytoplasm or perinuclear region, ruling out a role for HIF-1c in blocking AHA-1 nuclear translocation (Figure 5D). Instead, we observed a significant asymmetry in the nuclear distribution of Venus signal. HIF-1c/AHA-1 heterodimers clustered at the nuclear periphery in relatively DAPI-free regions, whereas HIF-1a/AHA-1 foci were dispersed throughout the nucleus and co-localized with DAPI (p<0.05; Figure 5E,F; Videos S3,4). Together, these results support a model in which HIF-1c acts as a dominant-negative regulator by binding AHA-1 in the nucleus, reducing the formation of transcriptionally competent HIF-1a/AHA-1 heterodimers and suggest that transcriptionally inactive HIF-1c/AHA-1 complexes may be selectively excluded from chromatin.

**FIGURE 5:**
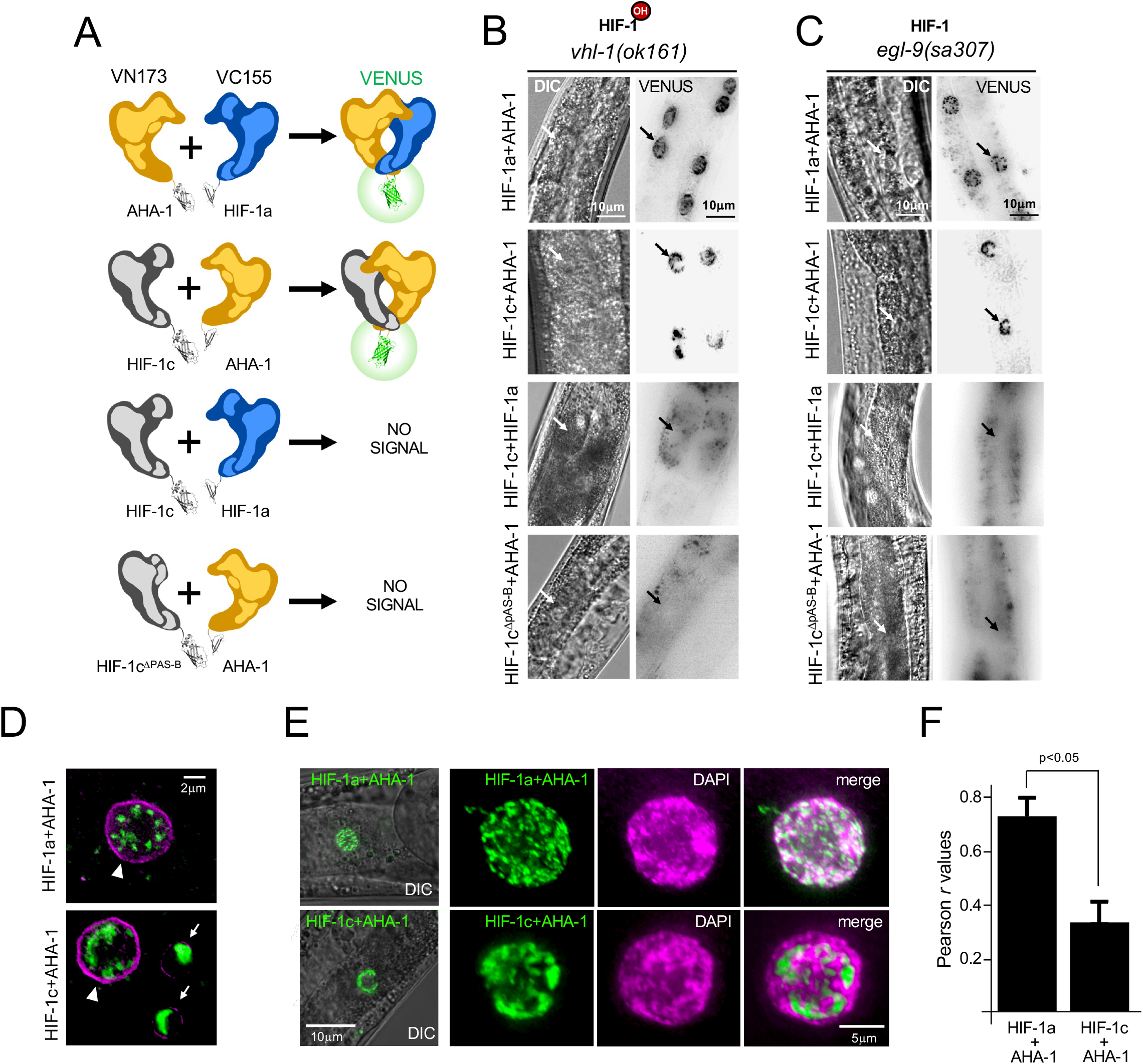
HIF-1c binds AHA-1 in the nucleus via the PAS-B domain. (A) BiFC interaction diagram; rows indicate tested combinations, with split-Venus VN173 constructs in the left column and VC155 constructs in the right; interaction outcomes (Venus signal) shown on the right. (B,C) DIC and inverted fluorescence images of intestinal cells in post-heat-shock *vhl-1(ok161)* (B) and *egl-9(sa307)* (C) worms expressing the BiFC constructs from (A); arrows mark intestinal nuclei. (D) Intestinal region of *vhl-1(ok161)* worms co-expressing AHA-1 + HIF-1a (top) or AHA-1 + HIF-1c (bottom) constructs and NPP-11::Scarlet; reconstituted Venus signal is observed in nuclei of intestinal (arrowhead) and hypodermal (arrow) cells. (E) DIC/BiFC Venus images of intact intestinal cells (left) and DAPI-stained nuclei of dissected intestines (right); native Venus fluorescence remains detectable after fixation. Top: AHA-1 + HIF-1a; bottom: AHA-1 + HIF-1c. (F) Quantification of DAPI/Venus co-localization (Pearson correlation coefficient) in dissected intestines (n=9 for HIF-1a + AHA-1, n=15 for HIF-1c + AHA-1).

### LIN-35 modulates *hif-1c* expression in neurons

To identify upstream regulators of *hif-1c* transcription in neurons, we exploited its reliance on the internal promoter. We screened a panel of mutants for neuronal transcription factors shown by ChIP-seq to bind intron 4 of *hif-1*, assessing the effect on expression of *Pi4::GFP* and *Phif-1::nRFP* reporters in a candidate-gene approach (Figure S7)^27^. Depletion of LIN-35, the *C. elegans* homolog of the Retinoblastoma protein (pRB), reduced neuronal *Pi4::GFP* expression without affecting canonical promoter activity (Figure 6A-C). This specificity of LIN-35 in regulating *hif-1c* prompted us to validate its role in HIF-1 signaling. Because *lin-35* mutants are known to exhibit enhanced somatic sensitivity to RNAi, which translates into ectopic silencing of GFP transgenes in the soma, we asked whether the reduced *Pi4::GFP* expression in LIN-35-depleted neurons might instead reflect increased RNAi^28^. To test this, we examined *Pi4::GFP* expression in *lin-35; mut-7* double mutants, in which RNAi is disabled by loss of MUT-7, an exoribonuclease required for siRNA biogenesis and RISC function. Indeed, *mut-7* mutations suppress the transgene-silencing phenotype of *lin-35* mutants^29^. RNAi-defective *lin-35; mut-7* worms still showed a significant reduction of *Pi4::GFP* expression, comparable to *lin-35* single mutants, whereas *Phif-1::nRFP* fluorescence was unaffected relative to *sasIs07* controls (p<0.001 and p=0.40, respectively; Figure 6D,E). Consistent with the reporter data, we found a significant reduction of endogenous *hif-1c* transcript levels in *lin-35(n745)* mutants by semi-quantitative RT-PCR (p<0.05; Figure 6F,G). Together, these results rule out a major role for the RNAi pathway in the loss of neuronal *Pi4::GFP* fluorescence in *lin-35* mutants and instead support a model in which LIN-35 directly promotes *hif-1c* transcription by mediating the activity of the internal promoter. If worms rely on LIN-35 to maintain physiological levels of HIF-1c, then depleting *lin-35* should elevate HIF-1-dependent gene expression, disrupt Eph signaling and cause neuronal defects similar to those seen in *vhl-1* and *hif-1(syb8674)* mutants. Indeed, *lin-35(n745)* worms show induction of *Pcysl-2::GFP* reporter (p<0.001), which is suppressed in a *lin-35; hif-1(ia04)* background (p=0.26), indicating elevated HIF-1 signaling in these animals (Figure 6H-J). This increase in HIF-1 activity parallels defects in HSN migration (24%; p<0.001; Figure 6K,L) and VC4/5 circuit formation (44%; p<0.001; Figure S8), supporting a critical role for LIN-35 in stabilizing neuronal HIF-1 signaling by regulating HIF-1c levels. Consistent with this model, MNs show robust expression of both *lin-35* and *hif-1c* (Figure 6M). LIN-35/pRB is a pocket protein that functions largely as part of a complex with DPL-1/DP1 and EFL-1/E2F, known as the DREAM (DRM) transcriptional complex^30^. Along with LIN-35, ChIP-seq data show that DPL-1 and EFL-1 pull down with intron 4 sequences, raising the possibility of DRM involvement in *hif-1c* regulation. However, we find that *Pi4::GFP* and *Phif-1::nRFP* expression levels were overtly unchanged in both *dpl-1(n3643)* and *efl-1(se1)* mutants, indicating LIN-35 is unlikely to regulate *hif-1c* transcription via DRM (Figure S9). In conclusion, analysis of candidate Pi4 transactivators revealed that LIN-35 positively regulates *hif-1c* transcription in neurons to maintain HIF-1 signaling at levels consistent with proper neuronal development.

**FIGURE 6:**
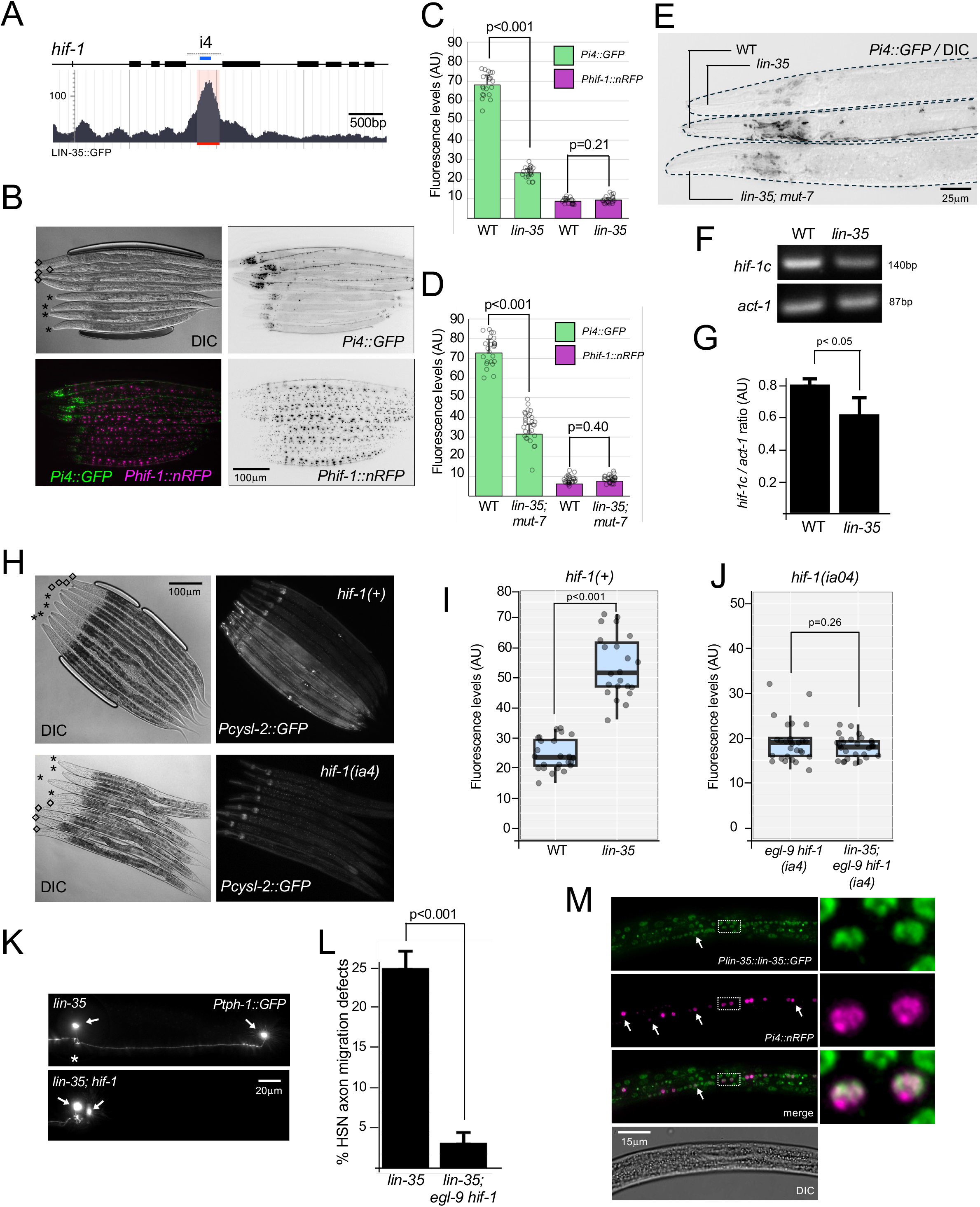
LIN-35 is a positive regulator of *hif-1c* transcription. (A-E) *hif-1c* expression is downregulated in *lin-35* mutants. (A) LIN-35 ChIP-seq read track across the *C. elegans hif-1* locus (Wormbase.org); the Pi4 region upstream of the putative TSS is marked by a blue bar, with the ChIP-seq peak within Pi4 marked by a red bar. (B) Representative images of *sasIs7* (diamonds) and *lin-35(n765); sasIs7* (asterisks) worms. (C) Fluorescence quantification of *Pi4::GFP* and *Phif-1*::*nRFP* in the nerve ring of *sasIs7* and *lin-35(n745); sasIs7* worms (n=24/genotype). (D) Fluorescence quantification of *Pi4::GFP* and *Phif-1*::*nRFP* in the nerve ring of *sasIs7* and *lin-35(n745); mut-7(pk720); sasIs7* worms (n=28/genotype). (E) DIC/GFP overlay images of *sasIs7*, *lin-35*, and *lin-35; mut-7* L4 worms expressing *Pi4::GFP*. (F) Semi-quantitative RT-PCR of *hif-1c* mRNA in whole-worm cDNA from WT (N2) and *lin-35(n745)* animals; relative *hif-1c* band intensities shown in (G), averaged across 4 independent experiments and normalized to *act-1* mRNA. (H-J) *lin-35* mutants have increased HIF-1 activity. (H) Worms expressing *Pcysl-2*::*GFP* (*nIs470*), with DIC images at left. Top two panels [*hif-1(+)*] show upregulated *Pcysl-2*::*GFP* in *lin-35(n745)* (asterisks) versus control *nIs470* (diamonds); this increase is abolished in a *hif-1(ia4)* background (bottom panels), with asterisks marking *egl-9(sa307) hif-1(ia4)* and diamonds marking *lin-35(n745); hif-1(ia4)*. (I,J) Quantification of *Pcysl-2*::*GFP* in the hyp7 syncytium (n=28) of *lin-35* mutants (I) and *lin-35; egl-9 hif-1* triple mutants (J). (K,L) HSN migration defects in *lin-35* worms. (K) *Ptph-1*::*GFP* images showing posterior HSN position (arrows) in *lin-35* mutants, quantified in (L). (M) Co-expression of *lin-35* and *hif-1c* reporters in MN nuclei (arrows) of an L3 larva; inset shows two MNs along the VNC.

## Discussion

### A novel HIF-1 inhibitor protein in *C. elegans*

Here we describe the first tissue-specific *C. elegans* HIF-1 isoform with dominant-negative function. HIF-1c restricts HIF-1 activity by limiting the nuclear availability of free AHA-1, thereby diminishing HIF-1-dependent transcription. This mechanism places HIF-1c among a broader family of dominant-negative HIF isoforms characterized in mice and humans, most of which arise as HIF-3α splice variants, including IPAS and HIF-3α4, respectively^31^. These isoforms, however, can inhibit HIF-1 through direct binding to HIF proteins ^7,26^. Instead, HIF-1c more closely resembles the mouse HIF-3α isoform NEPAS, which is expressed in the developing lung and heart, tissues in which sustained HIF-1 activity can cause hypoplasia and morphological defects, respectively^32^. Like HIF-1c, NEPAS inhibits HIF-1 indirectly by competing for a limiting pool of the shared dimerization partner ARNT^32^. A parallel strategy operates in the testis-specific variant hHIF-1αTe, which lacks the bHLH domain and instead sequesters ARNT into non-functional complexes, dampening HIF-1 signaling in the avascular, highly hypoxic testicular microenvironment^5^. In the testis, restraining stabilized HIF-1 is thought to prevent steroidogenic imbalances linked to male infertility^33^. Thus, like IPAS, the expression and function of *C. elegans* HIF-1c, mouse NEPAS, and human HIF-1αTe are each restricted to tissues particularly vulnerable to HIF-1-induced damage. Interestingly, tissue-restricted expression of hHIF-1αTe, a N-terminal truncated isoform like HIF-1c, is also conferred by an internal promoter within the *HIF1A* locus^5^. These isoforms diverge, however, in their sensitivity to oxygen: whereas IPAS and HIF-3α4 are oxygen-insensitive, NEPAS, hHIF-1αTe, and HIF-1c all retain oxygen-dependent stabilization, allowing these variants to exert maximal inhibition precisely when HIF-1 transcriptional activity peaks and the oxygen-dependent degradation machinery is least active^32^. Taken together, these parallels suggest that dominant-negative HIF variants represent a successful strategy adopted independently across metazoans to achieve context-specific calibration of HIF-1 activity.

### Safely wiring a nervous system under the threat of hypoxia

We propose that HIF-1c counterbalances the hypoxic response to control distinct aspects of neurodevelopment in *C. elegans* that would otherwise be disrupted by HIF-1 stabilization in the hypoxic embryo. These include failures in HSN migration, PVQ axon pathfinding errors, and VC4/5 axon outgrowth and branching defects. Surprisingly, the conserved oxygen-sensing machinery that degrades the bulk of HIF-1 is not sufficient on its own to protect neurons from HIF-1-mediated defects. Instead, our results indicate that competitive inhibition of HIF-1/AHA-1 complex formation by HIF-1c in the nucleus is a critical layer of HIF-1 signaling regulation during development. Previous studies implicated ephrin receptors as downstream mediators of the axon pathfinding and migration defects that arise upon HIF-1 stabilization in both *C. elegans* and zebrafish nervous systems^8,9^. The paradoxical effects of enhanced Eph signaling on nervous system wiring appear to represent a trade-off to ensure embryonic survival under hypoxia, as *vab-1* embryos fail to develop in low oxygen^8^. We have added VC4/5 to the roster of HIF-1-sensitive neurons along with DA/DB, PVQ, PVP and HSN and described defects in axon outgrowth and branching that disrupt formation of the egg-laying circuit and are similarly dependent on Eph signaling. Consistent with this, Eph has well-documented roles not only in cell migration and axon guidance but also in axon extension and branching^34–36^.

In which cells does HIF-1c act to control the fate of HSN, PVQ, and VC4/5 neurons? While HIF-1 is ubiquitously expressed, HIF-1c is predominantly produced in neurons, suggesting that its antagonism of HIF-1 signaling occurs primarily in these cells. In the most parsimonious model, HIF-1-dependent upregulation of VAB-1 in embryonic MNs titrates ephrins on the MN surface, effectively reducing available ligand to repel PVQ axons migrating along the VNC. This Eph barrier disruption model is supported by the observation that similar axon pathfinding errors occur in *vab-1* mutants, where Eph signaling is eliminated entirely^25^. A role for HIF-1c at the midline is consistent with its expression in MNs and is further supported by the VAB-1-dependent pathfinding defects of PVQ axons observed in *hif-1(syb8674)* mutants. However, because PVQ also expresses *hif-1c*, we cannot rule out a cell-autonomous function of HIF-1c in these neurons. In contrast, the protective effects of HIF-1c in developing HSN and VC4/5 neurons must be accomplished cell-non-autonomously, since these neurons do not express *hif-1c*. As with the PVQ axon pathfinding defects, pan-neuronal expression of HIF-1^P621A^ is sufficient to induce errors in HSN migration, consistent with a role for neuronal HIF-1c in this process^8^. During vulva organ formation in the L3 and L4 stages, the vulval epithelium plays a guidepost role in the innervation of the egg-laying circuit by restricting VC4/5 terminal arborization via expression of the neurexin-related BAM-1 and the N-glycanase PNG-1^22,23^. In addition, members of the PCP pathway PRKL-1, VANG-1, and DSH-1 which are expressed in VC neurons, HSNs and the vulval epithelium, control the number of VC4/5 neurites, preventing ectopic axons and ensuring a bipolar morphology^24^. HIF-1c-depleted mutants display both types of defects: increased branching and overextension of the terminal ends, as well as adoption of anterior-posterior (A-P) axis similar to V1/6 neurons often with supernumerary neurites running parallel to the VNC main tracks, resulting in a tripolar configuration. However, we do not detect *hif-1c* expression in the vulval epithelium or vulval muscle cells, suggesting that these non-neuronal tissues surrounding the developing vulva are not candidate sources of the extra VAB-1 generated by increased HIF-1 activity. Instead, we propose that upregulation of *vab-1* in *hif-1c*-expressing MNs proximal to the vulva may account for the disruption in Eph signaling underlying the defects in VC4/5. Correspondingly, eliminating Eph signaling in *vab-1* mutants results in VC4/5 wiring defects similar to those observed in upon depletion of *hif-1c* or *vhl-1*, though at a much lower penetrance, suggesting that HIF-1 stabilization may sensitize VC4/5 to ephrins. Alternatively, HIF-1 activity may disrupt Eph signaling through a gene other than *vab-1*. Indeed, ectopically expressing the Eph ligand EFN-2 in muscle can induce midline crossing of PVQ axons and the corresponding defects triggered by HIF-1 stabilization are partially suppressed in *efn-2* mutants^8^. It is possible, therefore, that VC4/5 defects in *hif-1c* mutants are the direct or indirect product of an Eph signaling imbalance at the ventral midline caused by upregulation of an Eph modulator gene in MNs.

### LIN-35/pRB controls HIF-1 signaling in neurons through HIF-1c

The dependence of *hif-1c* expression on Pi4 enabled us to dissect its transcriptional control and identify LIN-35 as new regulators of HIF-1 signaling in worms. LIN-35/pRB is a well-characterized cell cycle regulator that broadly remodels chromatin to repress genes required for mitotic entry while upregulating genes involved in differentiation and apoptosis^37^. Although less common, pRB can also act as a co-activator, directly upregulating genes at target promoters^38^. Similarly, in the *C. elegans* embryo, LIN-35 upregulates non-cell-cycle genes involved in neurogenesis independently of E2F^39^. Consistent with these precedents, our results suggest that LIN-35 associates with Pi4 via an unidentified, non-DRM factor to induce *hif-1c* transcription. This mechanism may parallel mammalian systems, where hypoxia modulates the phosphorylation-dependent cell cycle arrest function of pRB, a key aspect of its tumor-suppressor activity^40^, and where pRB physically interacts with the HIF-1 complex at HIF-1-responsive promoters to shape transcriptional outputs^41,42^. Together, these observations support a hierarchical model in which LIN-35 acts upstream of HIF-1 in neurons, inhibiting hypoxia-dependent gene expression by upregulating HIF-1c (Figure 7).

**FIGURE 7:**
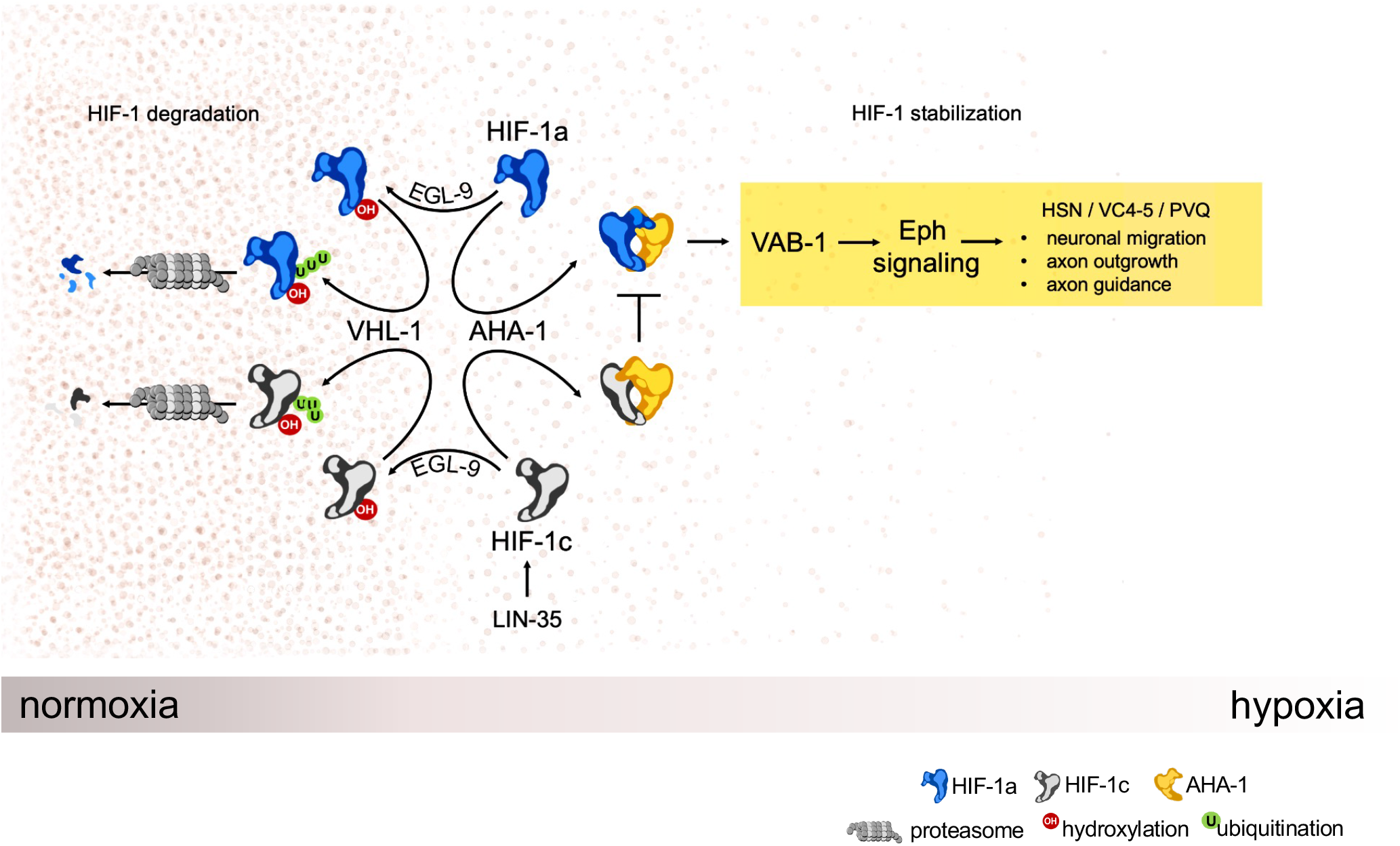
A model of HIF-1c-dependent of regulation of HIF-1 signaling in neurons.

### Implications to mammalian neurons

The neuronal defects resulting from HIF-1 activity in both worms and fish reveal a significant and conserved challenge to nervous systems across species, raising the question of its evolutionary significance to humans. Several studies implicate human HIF complexes in upregulating Eph signaling in cancer cells and endothelium, consistent with Eph acting as a downstream effector of HIF-1^43,44^. Moreover, hypoxia is a main driver of axon pathfinding errors in the CNS disruptions that are behind several neurodevelopmental disorders^10^. Although a direct link between excessive HIF-1 activity during mammalian brain development and connectivity defects remains unestablished, recent advances in non-invasive imaging that enable axon tracing and circuit-level resolution have revealed a correlation between periods of peak developmental hypoxia and the emergence of circuit disruptions. For example, laser-scanning photostimulation of the auditory cortex in rat models of hypoxia demonstrated hyperconnectivity in subplate circuits, the early scaffolding that connects thalamic inputs to the developing cortical plate^45^. In humans, neonatal hypoxia arising from preterm birth is known to damage subcortical circuits, a source of later cognitive impairment^46^. Thus, HIF-1 stabilization may also be behind the disruption in circuit formation in the developing human brain following hypoxic insults, underscoring a conserved need to shield neurons from sustained HIF-1 activity.

## Acknowledgments

We thank Dr. Alex Hajnal for kindly sharing the AH4533 strain. Some strains were sourced from the *Caenorhabditis* Genetics Center, funded by NIH Office of Research Infrastructure Programs (P40 OD010440). This work was funded by the Natural Sciences and Engineering Research Council of Canada (NSERC) grant 03751-2018.

## Author Contributions

ASA, CT, KC and CEC contributed with performing experiments. ASA and CEC contributed with designing experiments, writing and editing the manuscript.

## Declaration of Interests

The authors declare that they have no known competing financial interests or personal relationships that could have influenced the work reported in this paper.

## STAR★METHODS

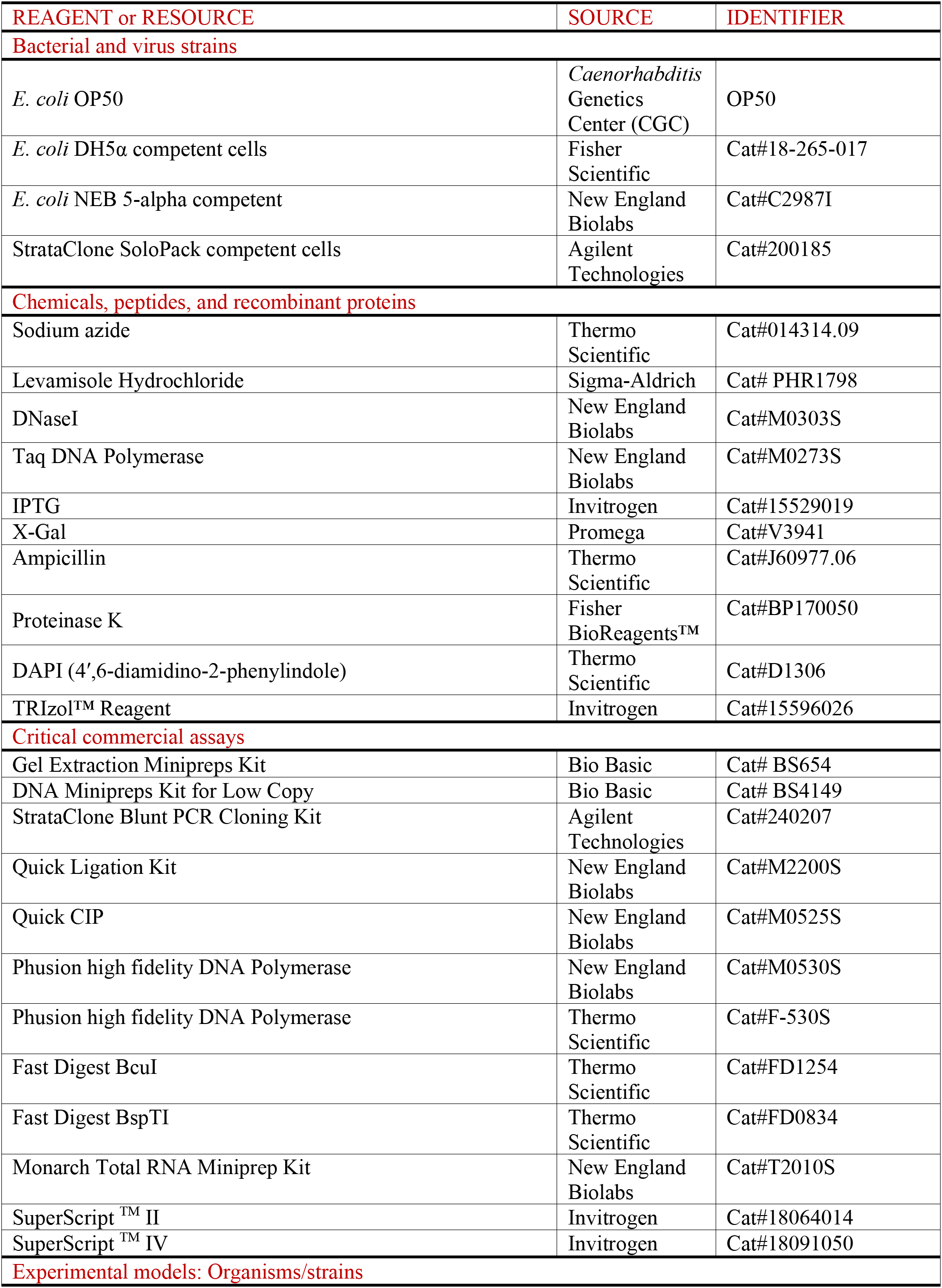
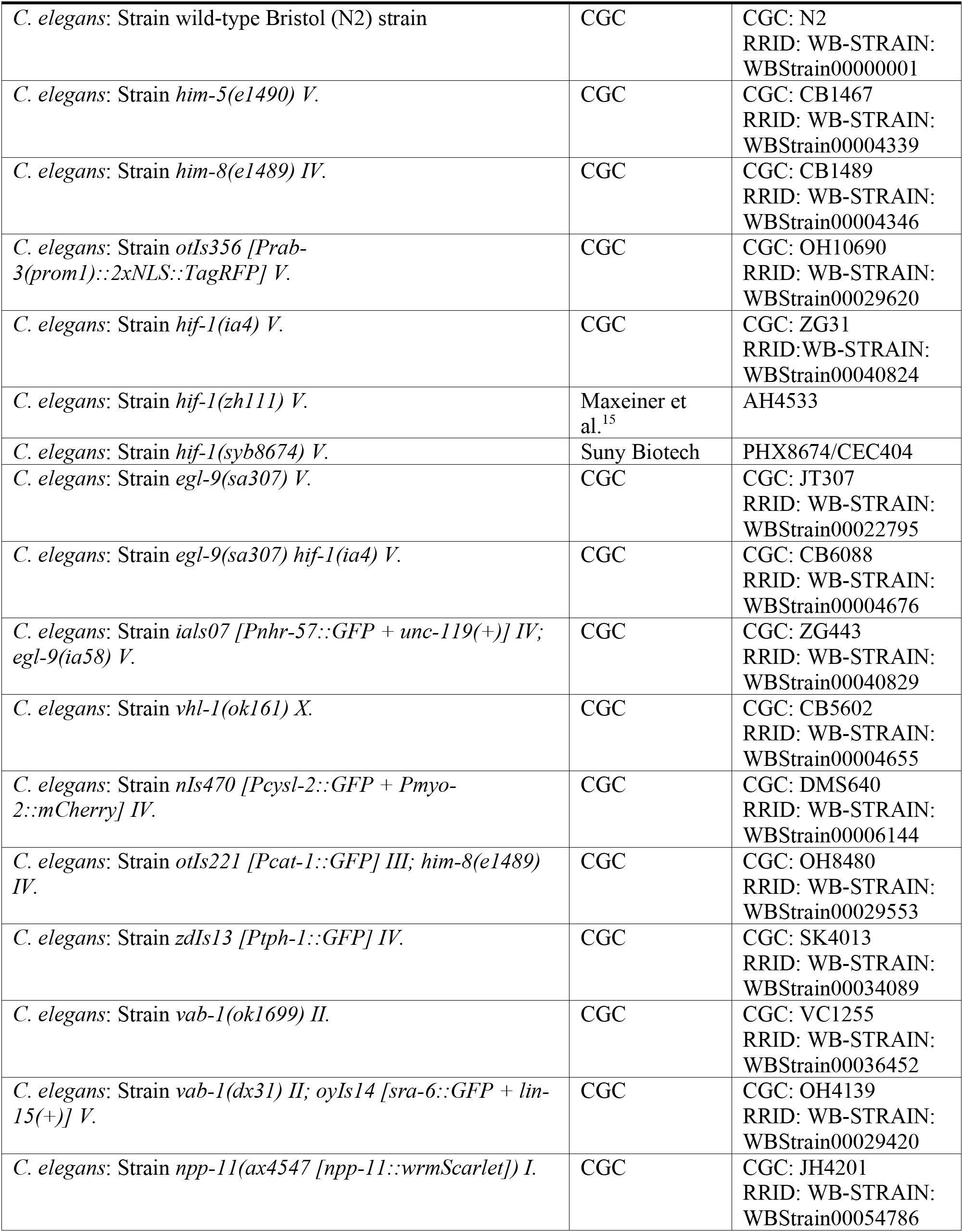
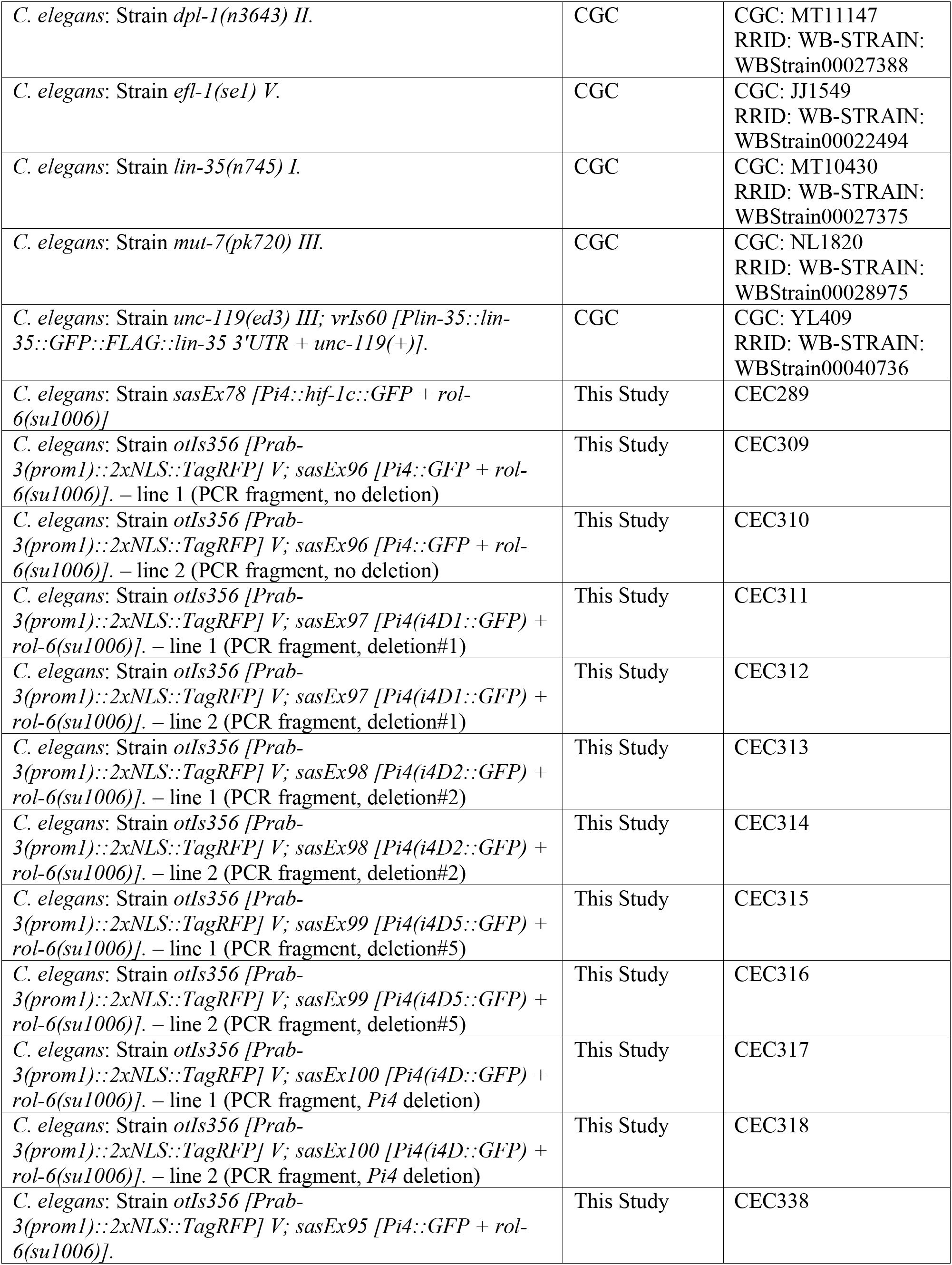
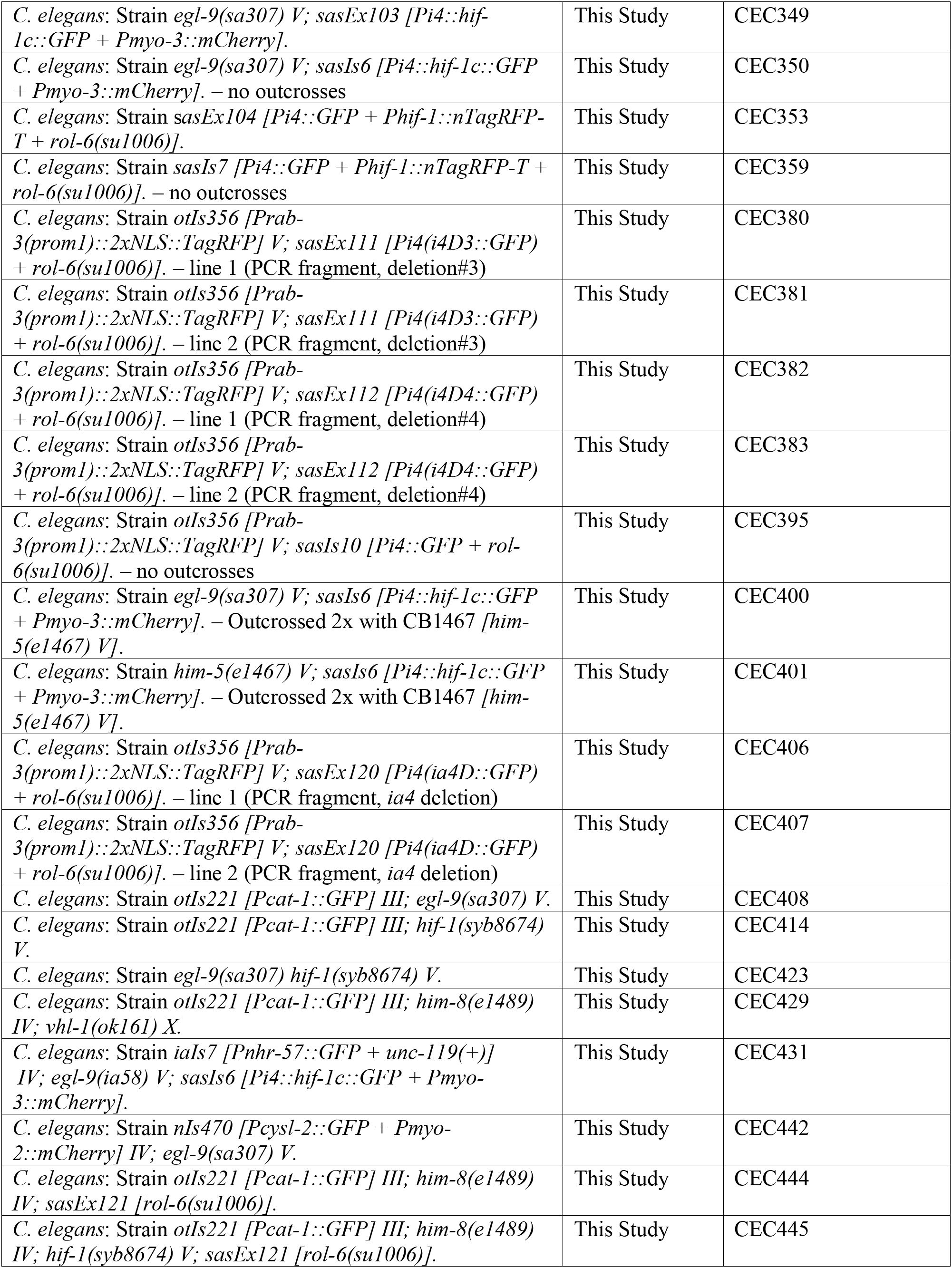
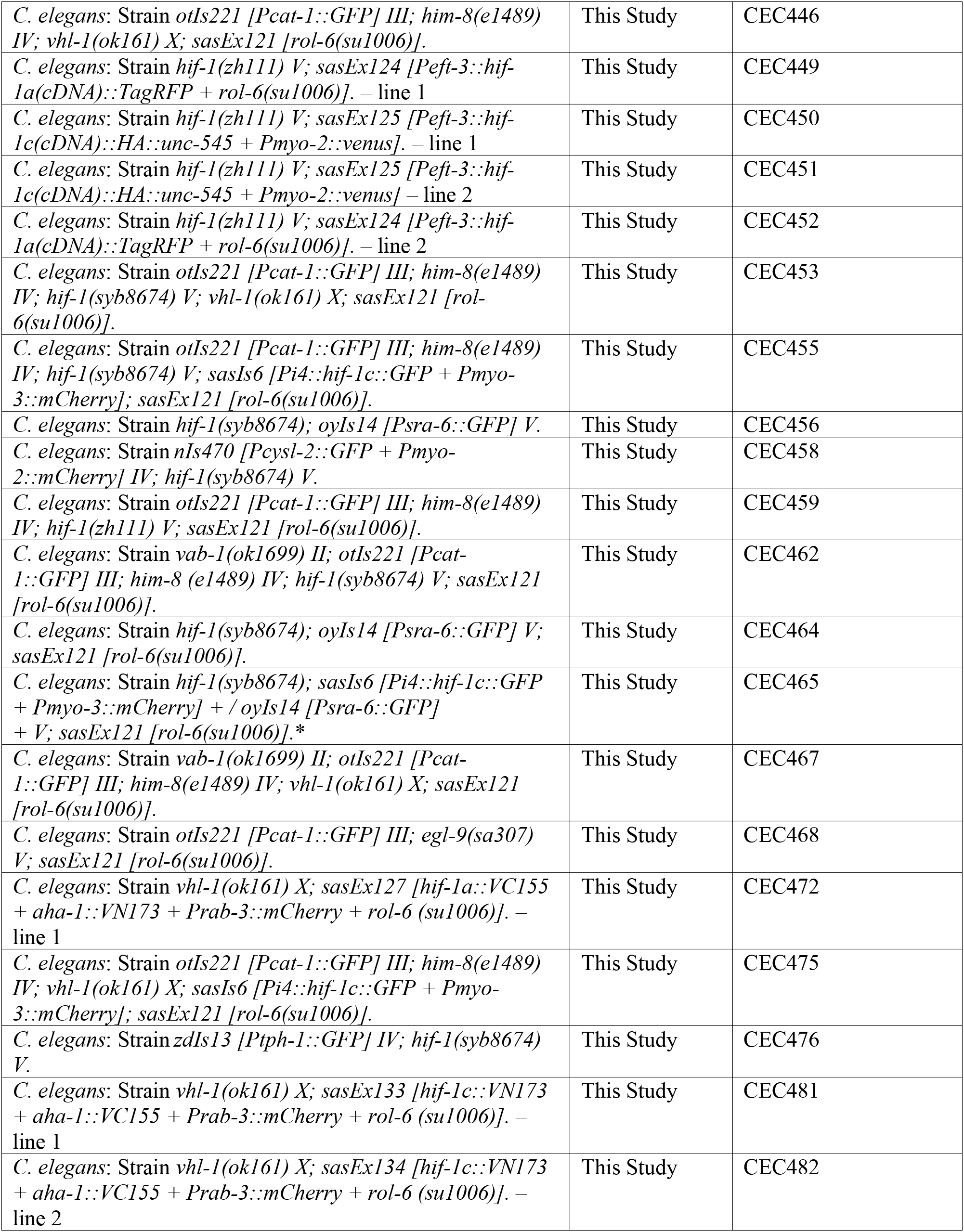
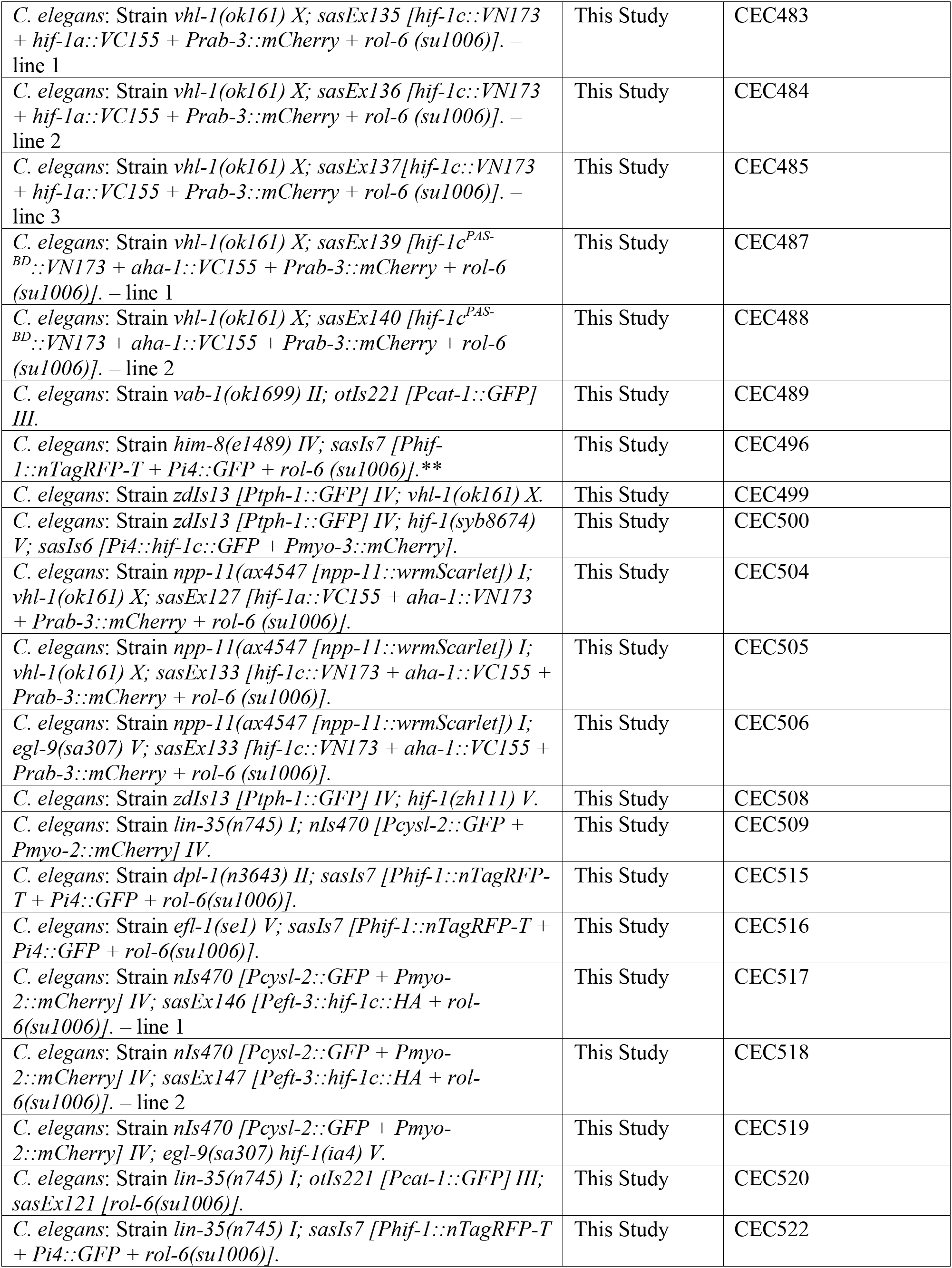
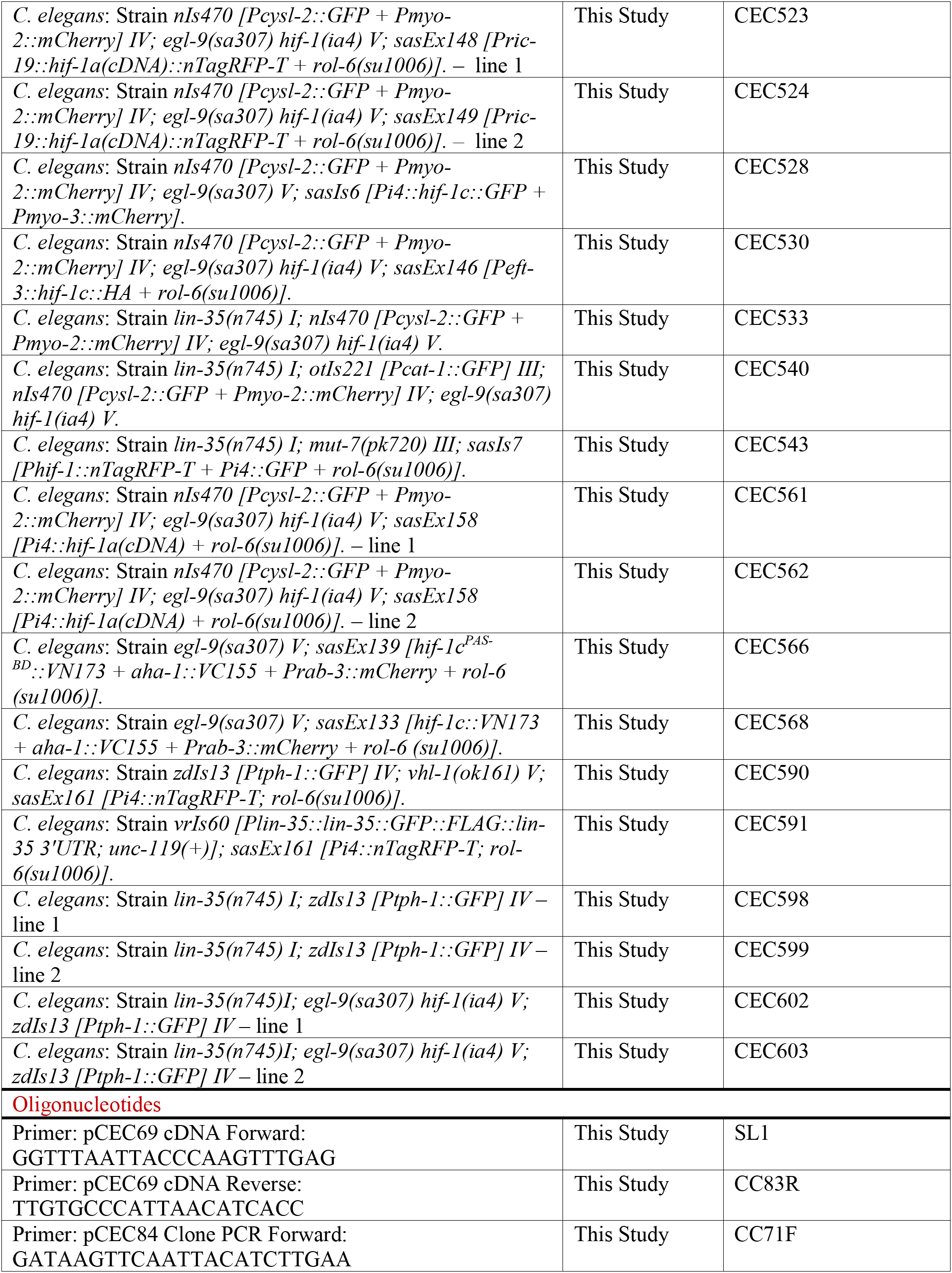
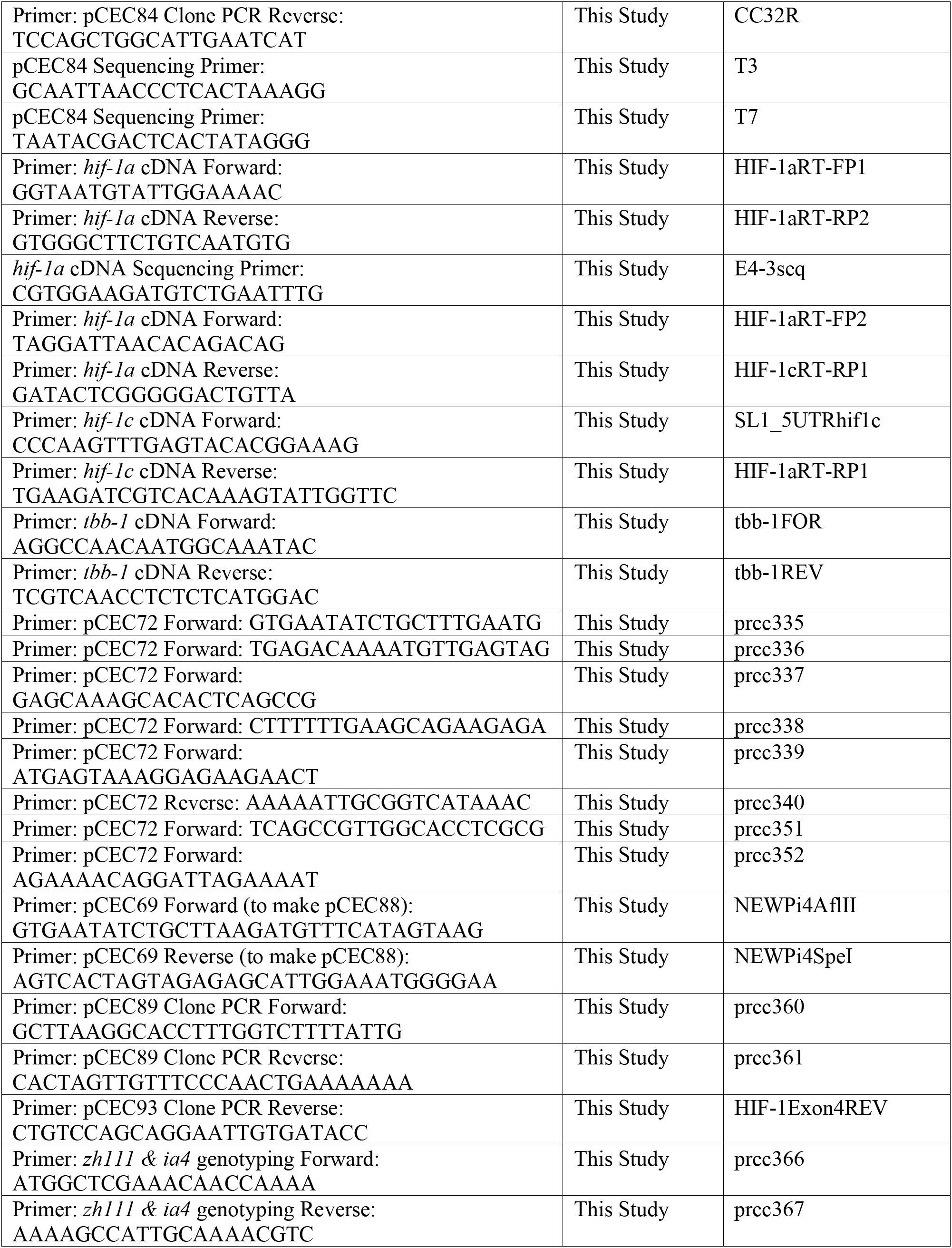
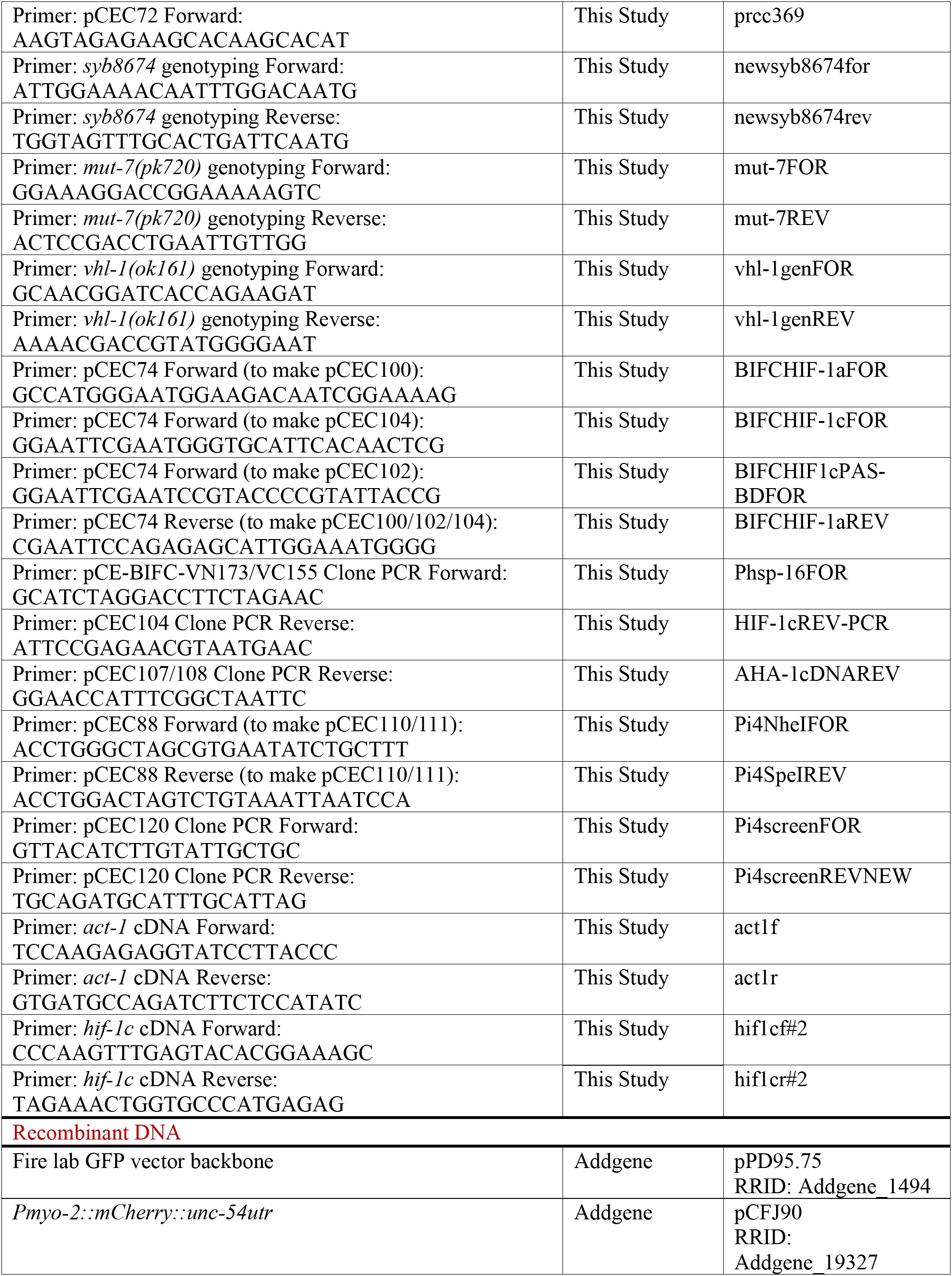
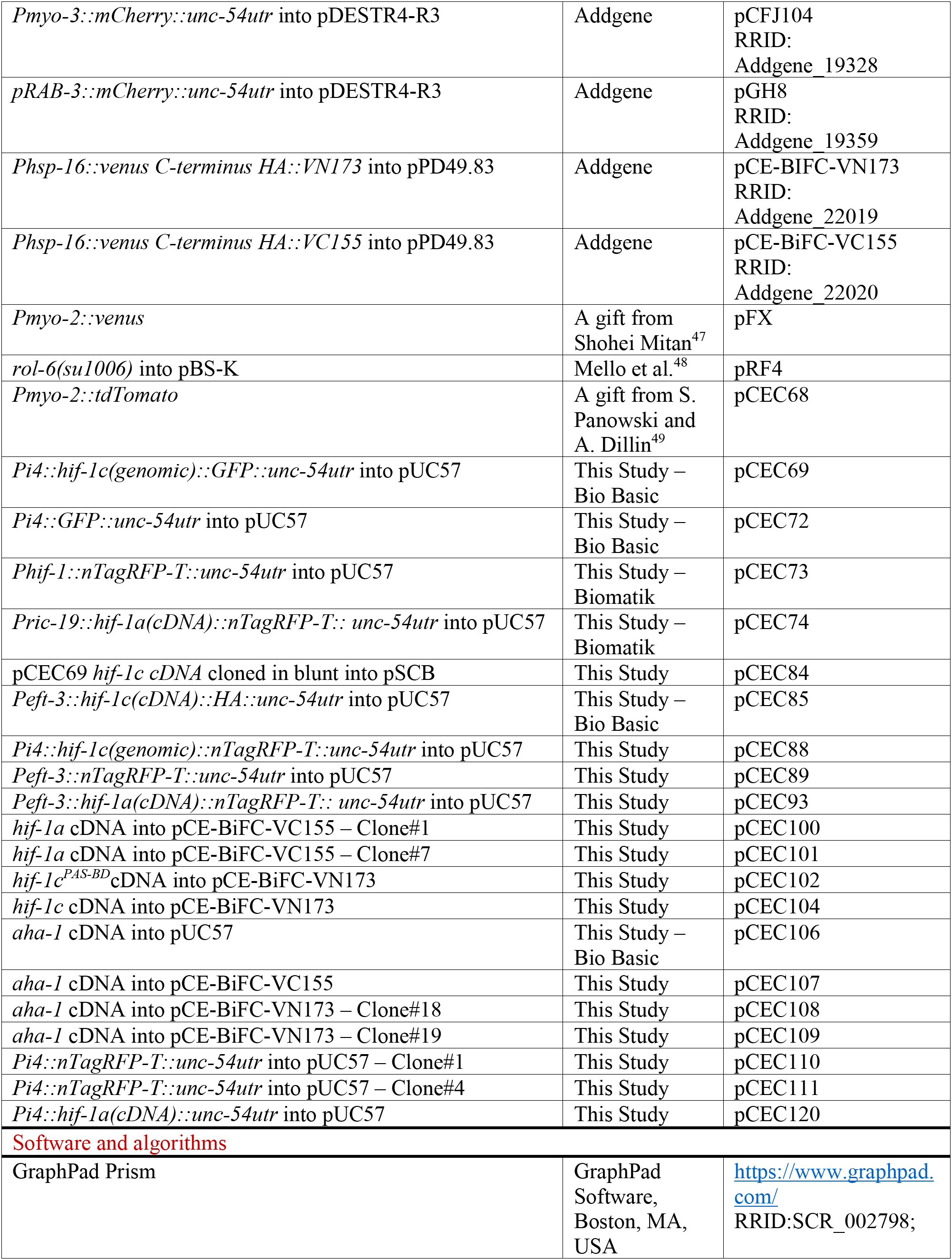
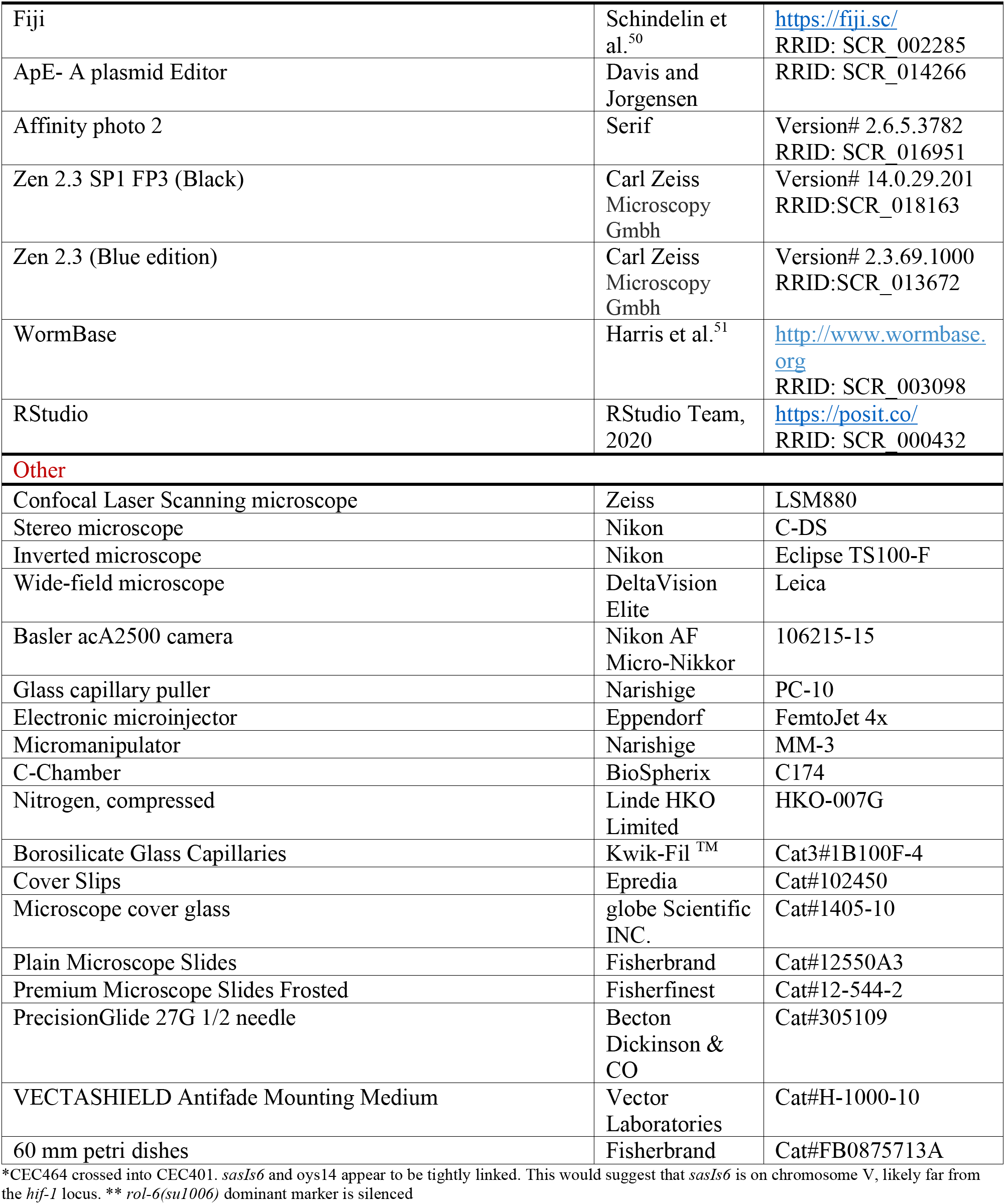

### Experimental model and study participant details

#### *C. elegans* husbandry and maintenance

*C. elegans* strains were maintained on nematode growth medium (NGM) agar plates seeded with *E. coli* OP50 at room temperature (∼20°C–22°C). The N2 Bristol strain served as the wild type^52^. Strains with temperature-sensitive (*ts*) alleles were kept at 16°C. For genetic crosses, 4 hermaphrodites were mixed with 8-10 males on a fresh 6 cm mating plate. Male worms for crossing were heat-shocked at 30°C for 4 hours. Progeny were then isolated, and target genotypes identified either by fluorescent marker visualization using a Nikon SMZ1500 stereo microscope or by single-worm PCR.

#### Constructs, strain generations, and transgenesis

*C. elegans* strains were obtained from the *Caenorhabditis* Genetics Center (CGC). The AH4533 [*hif-1(zh111)*] strain was kindly provided by Dr. Alex Hajnal. The CRISPR-engineered PHX8674/CEC404 [*hif-1(syb8674)*] strain was obtained from Sunny Biotech (Fuzhou, China). A complete list of strains used is provided in the Star Methods Table.

Transgenes were constructed using standard cloning techniques or Gibson assembly (Bio Basic). For PCR-generated inserts, Phusion high-fidelity DNA polymerase (Thermo Scientific) was used to amplify genomic or cDNA templates. Putative clones were screened by colony PCR with Taq polymerase to verify the presence and orientation of the insert, using primers listed in the Star Methods Table. The pUC57 vector (2710 bp, ampicillin resistance) served as the primary backbone for most cloning and subcloning procedures. A comprehensive list of all plasmids, either created (pCEC) or acquired from other laboratories, is provided in the Star Methods Table.

Transgenic *C. elegans* lines carrying extrachromosomal arrays were generated by germline microinjection into young adult hermaphrodites^53^. Injection mixtures contained plasmid DNA or gel-purified PCR-amplified DNA fragments at 20-50 ng/μL, along with pRF4 [*rol-6(su1006)*] or fluorescent markers such as *Pmyo-3::mCherry* as dominant marker. In this study, at least two independent, stable lines for each construct were isolated and characterized. Extrachromosomal arrays in transgenic strains used for behavioral experiments or tissue expression studies were integrated using UV irradiation. The resulting lines were then outcrossed twice with either the N2 strain or a high incidence of males (*him-5 or him-8*) background before proceeding with further analysis^54^.

#### Single Worm PCR genotyping

To genotype deletion alleles during strain construction, a PCR-based method was used. Briefly, worms from clonal F2 lines were isolated into 5 μL of fresh lysis buffer containing 0.2 mg of Proteinase K and kept on ice. After a 30-minute freeze/crack step at -80°C, the tubes were incubated in a PCR cycler for 90 minutes at 60°C to digest, then for 15 minutes at 95°C to inactivate Proteinase K. The lysed solutions served as templates in a 25 μL PCR with primers flanking the deletion site. The lysis buffer consisted of 50 mM KCl, 10 mM Tris (pH 8.2), 2.5 mM MgCl2, 0.45% NP-40, 0.45% Tween-20, 0.01% gelatin, and 0.2 mg/mL Proteinase K.

#### Confocal microscopy

A Zeiss LSM880 laser-scanning confocal microscope, equipped with GaAsP detectors and an Airyscan module, was used to capture DIC and fluorescence images. Worms were anesthetized in 15 mM sodium azide in M9 buffer, mounted on 2% agarose pads, and covered with a coverslip for live imaging. GFP fluorescence was excited with a 488 nm laser (2-5% power), and emission was collected at 493-550 nm. Nuclear TagRFP fluorescence was excited at 561 nm (3-7% laser power), and emission was collected at 570-650 nm. Sequential scanning was performed to prevent crosstalk. Images were captured as z-stacks using a Plan-Apochromat 40×/1.3 Oil DIC objective with a 0.5-1.0 µm step size in Zen Black (v2.3). Maximum-intensity projections and scale bars were generated in Zen Blue (v2.3). Unless otherwise noted, images are either single optical sections or maximum projections.

#### Widefield microscopy

A high-resolution deconvolution wide-field microscope (DeltaVision Elite, Leica) was used to image lines expressing functional HIF-1 transgenes (anesthetized with 10 mM Levamisol) or other transcriptional lines (anesthetized with 15 mM sodium azide). For fluorescence imaging, 0.2 μm optical sections were collected into z-stacks. Deconvolution of z-stack files was performed in Softworx (Applied Biosystems) using 10 cycles of aggressive ratio mode and a medium noise filter with a 200 nm cutoff. 3D reconstruction movies were also generated in Softworx. Unless otherwise noted, animals were synchronized and image at L4 stage. To study the developmental timeline of *hif-1a/c* expression, *sasIs7* homozygous late-stage embryos were collected in 10 mM Levamisol directly from NGM plates or from dissected from gravid worms for early-stage embryos. Embryonic stages were determined based on specific features, with at least 10 embryos imaged per stage. Dauer larvae were obtained from starved *sasIs7* plates.

#### Fluorescence quantification

For the promoter deletion analysis, fluorescence signals from GFP [*Pi4*-deleted constructs] and TagRFP [*Prab-3 (prom 1):: 2 xNLS:: TagRFP*] were systematically quantified from DVC interneuron cell bodies using a confocal laser-scanning microscope with identical imaging settings. DVC neurons are located near the tail neuron-poor region, just above the rectum^55^. GFP/RFP fluorescence ratios were calculated in Fiji/ImageJ (version 2.2.14.0.0) and normalized for array-specific effects. Control lines expressing the original *Pi4::GFP* (i4, positive control) and a GFP- only construct (i4Δ, negative control) were used to compare GFP/RFP ratios from the deletion constructs. Statistical significance was tested using unpaired t-tests.

Fluorescence quantification of *Pi4::hif-1c::GFP, Pcysl-2::GFP, Pnhr-57::GFP, Ptph-1::GFP, Phif-1::nTagRFP-T* and *Pi4::GFP* signals were performed on the DeltaVision system. Transcriptional reporters were imaged at 20x magnification. At this magnification, HIF-1c::GFP signals are not detectable. To minimize variability, age-matched control and tested L4 worms were aligned side by side on the same glass slide and imaged together at 20x or 40x magnification. At least 7 slides (28 worms per genotype) were imaged with consistent settings (exposure, binning, optical section). Z-stacks were deconvolved with identical parameters, collapsed, and saved as 8-bit grayscale images in Image J. With the exception of *Ptph-1::GFP*, fluorescence intensities were measured within a standardized pixel/area from all 8 worms per image, either from the nerve ring (between the two pharynx bulbs, for neuronal signals) or from the anterior end of the Hyp7 syncytium in the neck (for hypodermal signals), using Fiji’s measure function. Background fluorescence outside the sample area was subtracted afterward from each image set. Results are shown as average fluorescence (arbitrary units) from at least 7 slides, or as fluorescence ratios when applicable. Channel overlays and figure panels were prepared in Affinity Photo 2 (version 2.6.5).

To assess DAPI/Venus colocalization in BiFC-transgenic lines, Pearson correlation coefficients (*r*) were calculated for each nucleus using the Softworx co-localization module. A region around the intestinal cell nuclei was selected in deconvolved z-stacks of worms expressing HIF-1a+AHA-1 (n=9) and HIF-1c+AHA-1 (n=15). After setting a background threshold, signal intensities for each channel were measured in every optical section and compared pixel-by-pixel. High *r* values indicate strong overlap of signals across channels, whereas low values suggest the signals are not coincident.

#### Promoter deletion analysis

Staggered 5′-3′ deletion reporters of Pi4 were generated from the pCEC72 [*Pi4::GFP*] template using a PCR-based approach. Multiple 5′ primers targeting internal regions of intron 4 were paired with the same 3′ GFP primer to amplify *Pi4::GFP* constructs with Phusion Taq polymerase. PCR products were gel-extracted, and their concentrations were measured with a NanoDrop spectrophotometer. Eight constructs covering intron 4: i4, i4Δ1, i4Δ2, i4Δ3, i4Δ4, i4Δ5 i4Δ and *ia4*Δ were generated. These constructs were expressed as stable extrachromosomal arrays in a*Prab-3 (prom 1)::2xNLS::TagRFP* (OH10690) genetic background that marks all neuronal nuclei. Two independent transgenic lines for each construct were used for quantification. Control strains CEC309/CEC310 expressing full-length intron 4 in *Pi4::GFP* (*i4*) and negative control strainss CEC317/CEC318 expressing only GFP sequences (no intron 4, i4D) were used to establish expression thresholds.

#### cDNA analysis

To confirm the identity of the transcript produced by *Pi4::hif-1c::GFP*, a 5’ Rapid Amplification of cDNA Ends (RACE) approach was employed. Total RNA was extracted from a mixed population of *Pi4::hif-1c::GFP* expressing worms using TRIzol®. The RNA concentration was determined using a NanoDrop spectrophotometer and first-strand cDNA synthesis was carried out with SuperScript™ II (Invitrogen) using Oligo(dT)12–18 primers. The following RT-PCR utilized Phusion High-Fidelity DNA polymerase (NEB) with SL1 as the forward primer and CC83R as the 3’ GFP-specific primer. The resulting cDNA fragment was gel-extracted and blunt-end cloned into competent cells via the StrataClone Blunt PCR Cloning Kit (Strategene). Colony PCR screening was performed with standard Taq polymerase and primers listed in the Star Methods Table. Plasmids were sequenced using T3/T7 primers at the NRC-Saskatoon sequencing facility. cDNA of N2, *hif-1*(*syb8674)* and *hif-1(zh111)* worms were produced as described above and PCR reactions using primers specific for *hif-1a* and *hif-1c* used to determine the impact of *zh111* and *syb8674* in expression of *hif-1a* and *hif-1c*. Tubulin (*tbb-1*) cDNA amplification was used as a loading control (see the Star Methods table for primers).

#### Hypoxia Sensitive Assay (HSA)

Survival under hypoxia was tested in a c-chamber connected to a calibrated ProOx P110 O_2_ regulator (Biospheryx). Hypoxia was achieved by continuous infusion of pure (>99.95%) nitrogen gas (Linde HKO Limited). An additional portable O_2_ sensor was placed inside the chamber to ensure proper atmospheric conditions during incubations. Tests were conducted at room temperature. A 200 mL beaker with distilled water was added to maintain constant humidity.

HSA test plates were prepared by depositing 30 μL of OP50 liquid broth (OD600 = 1-3) onto the center of an NGM plate using a P100 pipette to form a circular lawn approximately 0.5 cm in diameter. NGM plates from the same batch were used to avoid subtle differences in agar composition and humidity. The bacterial lawns were incubated for 2 days at room temperature before use.

Each experiment tested one plate per genotype, with three hypoxia and three normoxia trials for each genotype. On day 1, 40-50 L4-stage worms of the same genotype were transferred to fresh NGM/OP50 plates. After 24 hours (day 2), young adults were placed on a glass slide with 35 μL of M9 buffer and dissected near the vulvar region using hypodermal needles to release embryos. The embryo suspension was gently pipetted to free the embryos from the carcasses. Under a stereoscope, two- to four-cell embryos were identified and aspirated with a mouth pipettor attached to a glass capillary, avoiding carcasses, bacteria, older embryos, or hatched larvae that could clog the capillary or interfere with the experiment. The embryo-containing M9 solution was pipetted onto a spot just outside the bacterial lawn, marked with a Sharpie. This was repeated until about 35-50 embryos were collected. Immediately after the solution dried, the number of embryos in the marked area was counted and recorded, with any older embryos or larvae removed. Plates were then placed inside the hypoxia chamber. Once all plates were inside (lid face up for easy gas exchange), the chamber was sealed. O_2_ levels stabilized at 0.03% after 30-40 minutes. Given the 30-minute dissection, most embryos were likely at the 8-cell stage when the hypoxia treatment started. Plates were incubated for 96-hour incubation with daily O_2_ level checks. Experiments with accidental reoxygenation were excluded from analysis. At the end of incubation time, the system was shut down, plates were taken out of the hypoxic chamber and immediately scored for the number of live L4/adult worms. Viability was determined by dividing the number of viable worms observed by the total number of embryos recorded for the respective plate. Control assays conducted in room air were performed separately to evaluate viability under normoxic conditions. Data represent the mean of three independent experiments for each genotype.

#### Egg-laying Assay

The number of eggs laid by synchronized young adult worms during two consecutive periods (0-2 and 2-4 hours) were scored. At least 10 synchronized L4 worms per genotype were transferred onto fresh NGM agar plates seeded with *E. coli* OP50 and incubated at 20°C for 24 hours before testing. Test plates were prepared by spreading bacteria with a glass spreader to create a 2 cm² bacterial lawn at the center. Lawn thickness was kept consistent by incubating plates for two days at room temperature. On day 1, adult worms were placed individually onto test plates and transferred to new plates after 2 hours. Eggs laid during the first period (0-2 hours) were counted directly on the original plates. After 4 hours, worms were removed, and eggs on the second plates (2-4 hours) were counted. Results are shown as the average number of eggs laid over 4 hours. Three replicates per genotype were conducted.

#### Neuronal defects quantification

Neuronal defects in cell migration, polarity, axon outgrowth/guidance and abnormal branching were evaluated using lines with GFP reporters that highlight a subset of neurons (see Star Methods Table). The rescue of *hif-1c* neuronal defects was assessed in *hif-1(syb8674)* backgrounds expressing a *hif-1c* translational reporter driven by the native *Pi4* promoter (*sasIs6* allele).

*Psra-6::GFP* (*oyIs14* allele)-expressing L2 worms were examined to analyze PVQ axon structure along the VNC. Typically, PVQ axons run in separate, parallel paths within the two VNC fascicles, except in a short proximal region near the cell body, where they briefly come into proximity. Abnormal guidance was identified when the axons of two PVQ neurons temporarily followed the same track over short distances (partial midline crossing) or when they followed the same track from the tail to the head ganglia after crossing (full midline crossing). Three independent experiments were performed with at least 130 worms per genotype in each experiment.

Adult synchronized *Ptph-1::GFP*-expressing worms (*zdIs13*) were used to assess axon guidance and migration defects in the two HSN neurons. In these experiments, two HSN defects were detected: a migration defect in which the cell body of one HSN fails to migrate to the midsection and retains a posterior position in the adult worm, and axon midline crossing defects in which HSN axons fail to remain on their respective tracks and cross the midline anteriorly toward the head. For quantification of HSN migration defects, only worms with at least one HSN cell body located 100 μm or more from the vulva slit were scored as defective. An average of n = 50 worms per genotype was scored across three independent experiments. Data are presented as the average of the three experiments. Of note, *zdIs13* expression in the head ASG neurons is sensitive to HIF-1^17^ and, in this context, also served as a neuronal reporter for HIF-1 activity.

The *otIs221* transgene (*Pcat-1::GFP*) was employed to study VC4/5 neuron morphology. Synchronized day 1 adult worms expressing *Pcat-1::GFP* were examined for defects in VC4/5 neurons. A *rol-6*-expressing array (CEC444 line) was introduced via microinjection to maximize ventral vulva positioning on slides. Only worms in which a ventral view of the egg-laying circuit was observable were scored. Three defect classes were identified: (i) VC4 or VC5 with A-P polarity where one axon fails to travel around the vulva; (ii) VC4 or VC5 with an extra process running parallel to the VNC (tripolar configuration) and (iii) over-extension of terminal branches or extra neurites often crossing the midline. Defected were scored and pooled together, with 60 worms per genotype in each experiment. Averages across three experiments are presented.

#### Bimolecular Fluorescence Complementation (BiFC) assay

For BiFC interactions, an *aha-1* cDNA *kpnI* fragment was subcloned from pCEC106 into the *Phsp-16* BiFC vectors pCE-VC155 and pCE-VN173. PCR-amplified cDNA fragments for *hif-1a* (NcoI/EcoRI) and *hif-1c* (EcoRI) were cloned in frame with pCE-VC155 and pCE-VN173^56^, respectively. A mutated *hif-1c* PCR amplicon missing the PAS-B domain (HIF-1c^PAS-BΔ^) was generated by using the 3’ primer BIFCHIF1cPAS-BDFOR and cloned into the EcoRI site of pCE-VN173. After directional check by clone PCR and sequencing, constructs were co-injected into *vhl-1(ok1621)* worms along with the *rol-6* marker plasmid (pRF4) and *Prab-3::mCherry* (pGH8) as described above. Venus complementation tests involved combinations of HIF-1a+AHA-1; HIF-1c+AHA-1; HIF-1a+HIF-1c; and HIF-1c^PAS-BΔ^+AHA-1. For each pairwise combination, two independent lines were established and maintained at 16°C. To assess complementation, 50 L4 rollers per line were placed on NGM-OP50 plates for 20 hours at 16°C, then heat-shocked at 33°C for 3 hours. After a 1-hour recovery at 16°C, worms were mounted in 10 mM Levamisole in M9 and immediately imaged. A few non-heat-shocked worms were checked to confirm heat induction of *Phsp-16* ^57^. Subcellular localization of Venus foci was examined using transgenic arrays with complementary BiFC constructs crossed into worms expressing the nuclear envelope marker *npp-11* (*npp-11::Scarlet; ax4547* allele). Hermaphrodites from one line expressing HIF-1a+AHA-1 and HIF-1c+AHA-1 were crossed into the *egl-9(sa307)* mutant to transfer the arrays. After heat shock, the lines were evaluated to determine whether non-hydroxylated HIF-1a/HIF-1c accumulation in the *egl-9* background affected their association with AHA-1.

For DAPI staining, heat-shocked worms were dissected on a glass coverslip in 30 μL of M9, with the vulva sliced using hypodermal needles. The dissected worms were then immediately transferred to a lysine-coated glass slide (Fisher Scientific) and flipped onto a -80°C chilled metal block to snap freeze. The coverslip was carefully pried off with a razor blade, allowing dissection remnants like intestines, embryos, and gonads to adhere to the positively charged slide surface. The slides were fixed promptly in pre-chilled (-20°C) 100% methanol for one minute, washed three times with PBS-T, then stained with 5 μg/mL DAPI in PBS-T for 10 minutes at room temperature. After a 30-minute PBS-T wash, the slides were mounted with 10 μL of Vectashield® mounting medium. This methanol fixation preserves native BiFC Venus fluorescence while permitting DAPI to access DNA. Z-stacks of 0.2 μm optical sections from intestinal cells in the DAPI and FITC channels were collected on the DeltaVision system and deconvolved with identical settings. To assess Venus expression in neurons, cells co-expressing *Prab-3::mCherry* in the head ganglia were identified and examined for Venus fluorescence. The 1X PBS-T solution (pH 7.4) contains: 137 mM NaCl, 2.7 mM KCl, 10 mM Na_2_HPO_4_, 1.8 mM KH_2_PO_4_, and 0.1% Tween 20.

#### Gene candidate screen for *Pi4* regulators

To identify regulators of *hif-1c* expression, the *C. elegans* modENCODE database for neuronal GFP::TFs ^27,58–60^ was analyzed for binding sites overlapping with the internal promoter region in intron 4 (V:20782386-20782529). Eight such TFs showed statistically enriched read values in the Pi4 region. The corresponding viable homozygous mutant strains were sourced from CGC and crossed into a *sasIs7* background to evaluate *Pi4::GFP* and *Phif-1::RFP-T* expression levels as compared to *sasIs7* controls (see Star Methods Table).

#### Semi-quantitative RT-PCR

Semi-quantitative RT-PCR was performed to assess *hif-1c* expression levels in *lin-35* mutants versus N2 worms. *lin-35(n745)* RNA was isolated from animals cultured at 16°C. Total RNA was purified from TRIzol®-extracted samples using the Monarch® Total RNA Miniprep Kit (NEB) according to the manufacturer’s instructions. DNase I treatment was applied to eliminate genomic DNA contamination. cDNA synthesis used 250 ng of total RNA. RT-PCR was performed with primers listed in the Star Methods Table for *hif-1*-specific amplicons and *act-1* (actin) control. Four paired reactions for *hif-1c* and *act-1* were conducted with *lin-35* and N2 cDNA templates under identical conditions. PCR products were run together on the same gel and imaged. Band densities for *hif-1* and *act-1* were quantified using Fiji. The normalized *hif-1c/act-1* ratio for each genotype and experiment was calculated and averaged across four replicates.

#### Statistical Analysis

We performed statistical analyses using GraphPad Prism, RStudio, or online tools. Data are shown as mean ± standard deviation (SD) unless specified otherwise. For related groups, paired two-tailed Student’s t-tests were applied; for independent groups, unpaired two-tailed Student’s t-tests assuming equal variances were used. A p-value < 0.05 was deemed statistically significant.

## Supplemental Figure Legends

**FIGURE S1:**
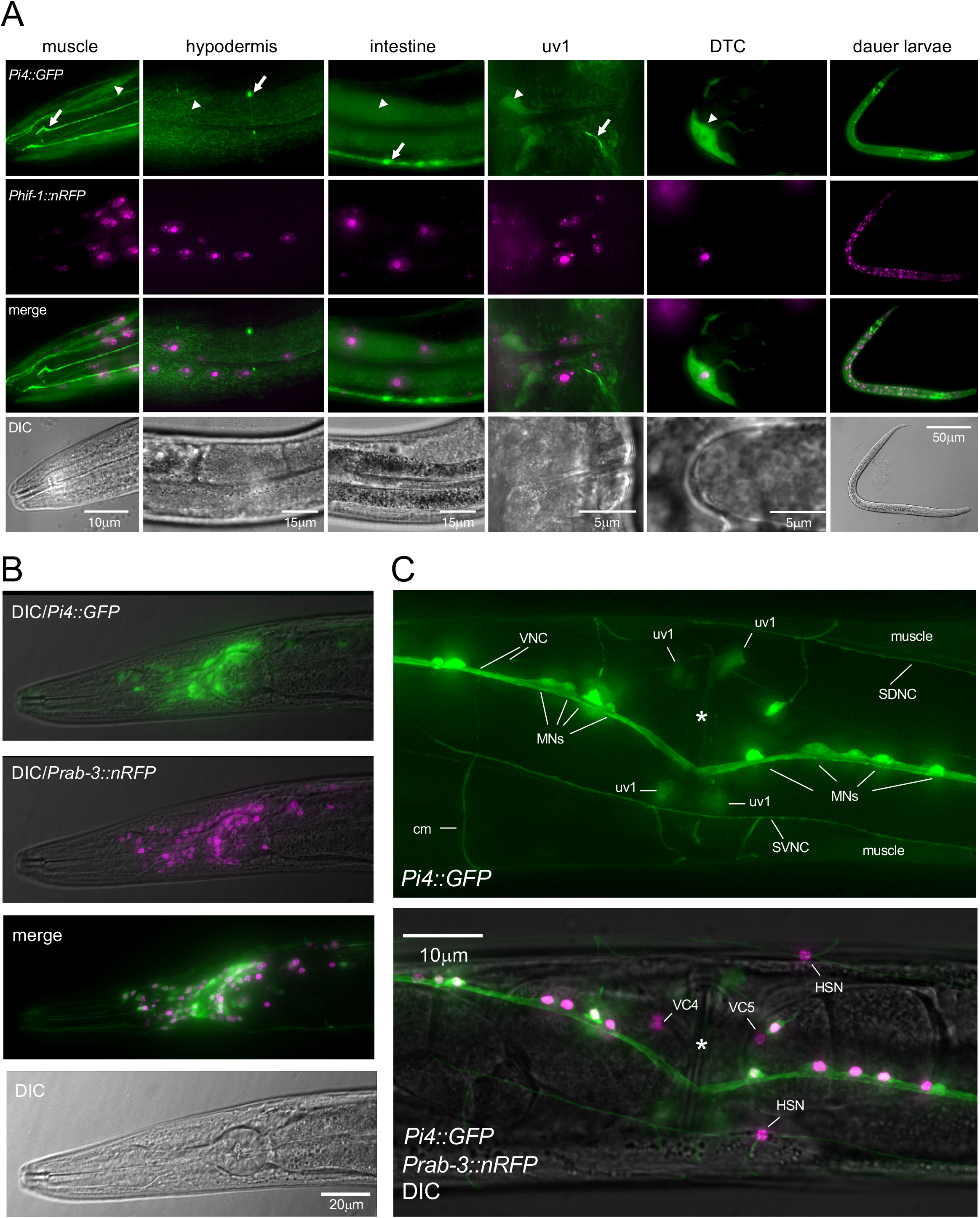
*Pi4::GFP and Phif-1::nRFP* expression in non-neuronal and neuronal cells. (A) *Pi4::GFP* signal is strongly detected in the DTC (distal tip cell of the somatic gonad) and weakly in head muscle cells, hypodermis, intestine and uv1 cells. All tissues with detectable *Pi4::GFP* signal also express *Phif-1::nRFP*. Compare expression levels in neurons (arrows) with those in weakly expressing non-neuronal tissues/cells (arrowheads). (B) Co-localization of *Pi4::GFP* and *Prab-3::nRFP* in head neurons of an adult worm. (C) *hif-1c* expression in VNC neurons. A ventral view of the vulval region in an adult worm co-expressing *Pi4::GFP* and *Prab-3::nRFP.* Robust *Pi4::GFP* expression is observed in MNs, the VNC and the uv1 cells, but is notably absent in VC4/5 and HSNs. SDNC – subdorsal nerve cord; SVNC – subventral nerve cord; cm – commissure.

**FIGURE S2:**
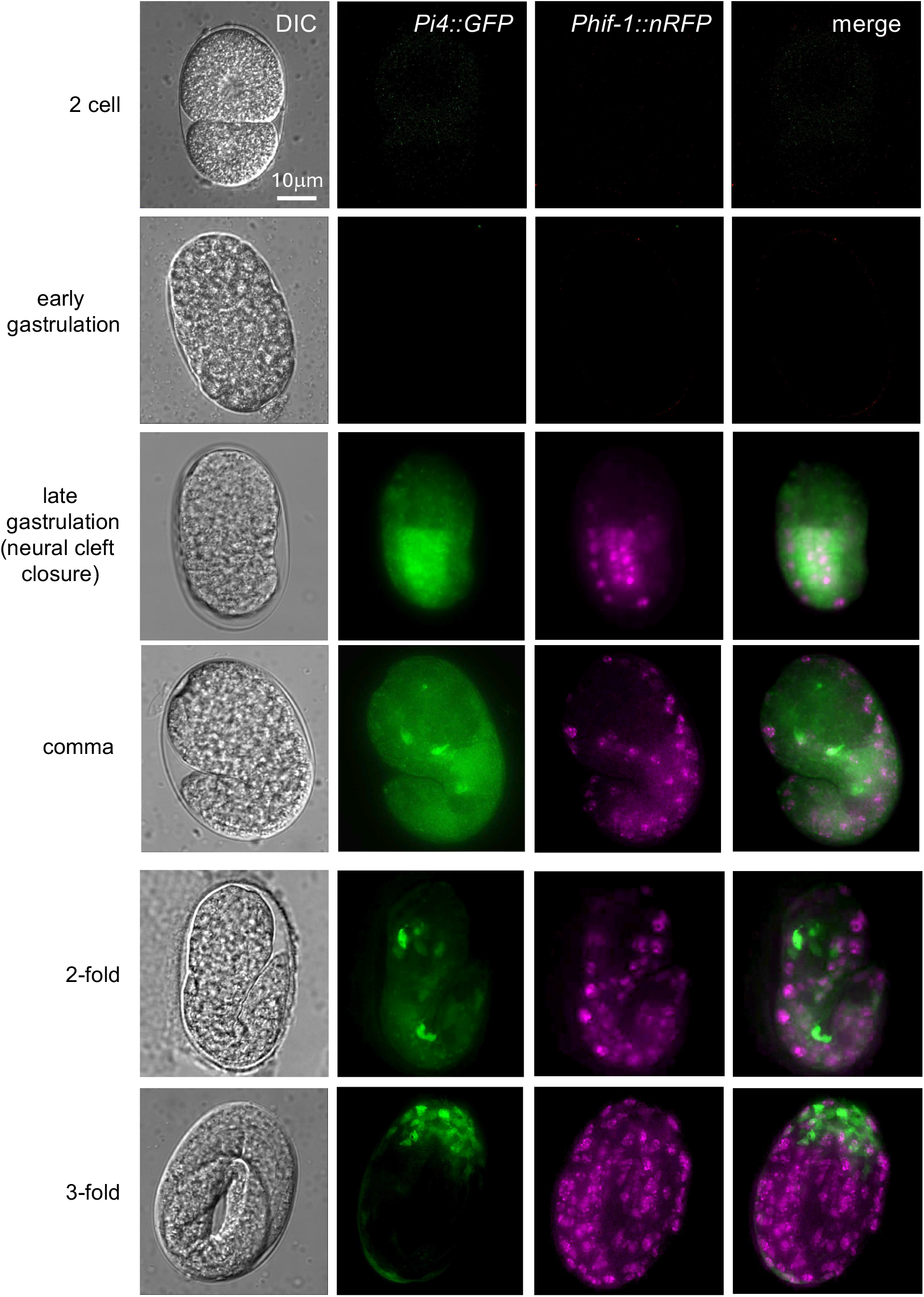
Expression of *Pi4::GFP* and *Phif-1::nRFP* during embryonic development. Expression of both reporters first becomes detectable during late gastrulation, when *Pi4::GFP* and *Phif-1::nRFP* signals are observed in neuroblasts. As embryos develop, *Pi4::GFP* is readily observed in the head and tail ganglia whereas *Phif-1::nRFP* expression expands beyond the nervous system.

**FIGURE S3:**
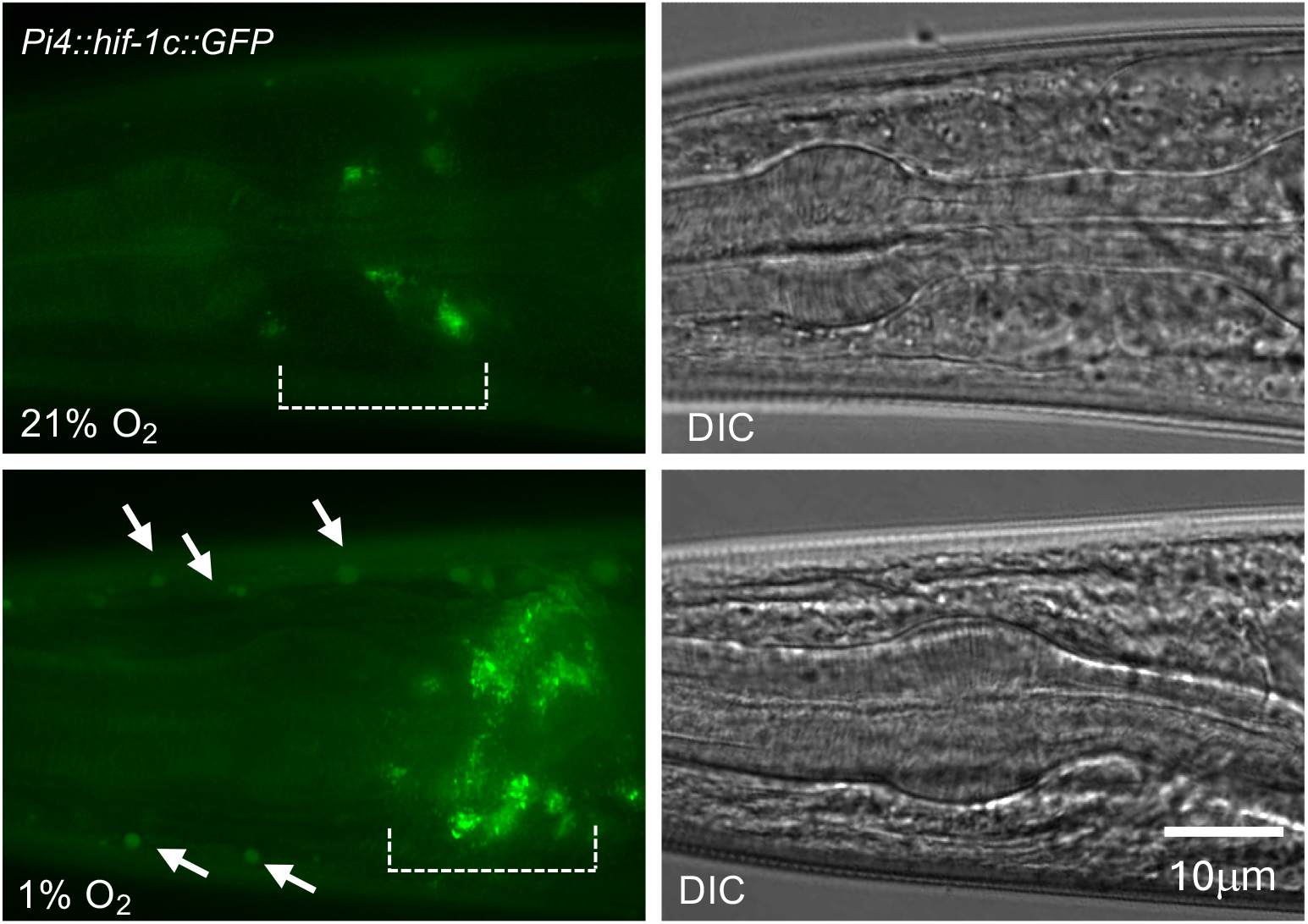
Oxygen sensitivity of HIF-1c. HIF-1c::GFP stability in anterior ganglia neurons (marked by a traced line) in the heads of L4 worms raised in room air (21% O_2_) or under hypoxia (1% O_2_). Arrows indicate a weak signal in nuclei of head muscle cells grown under hypoxic conditions.

**FIGURE S4:**
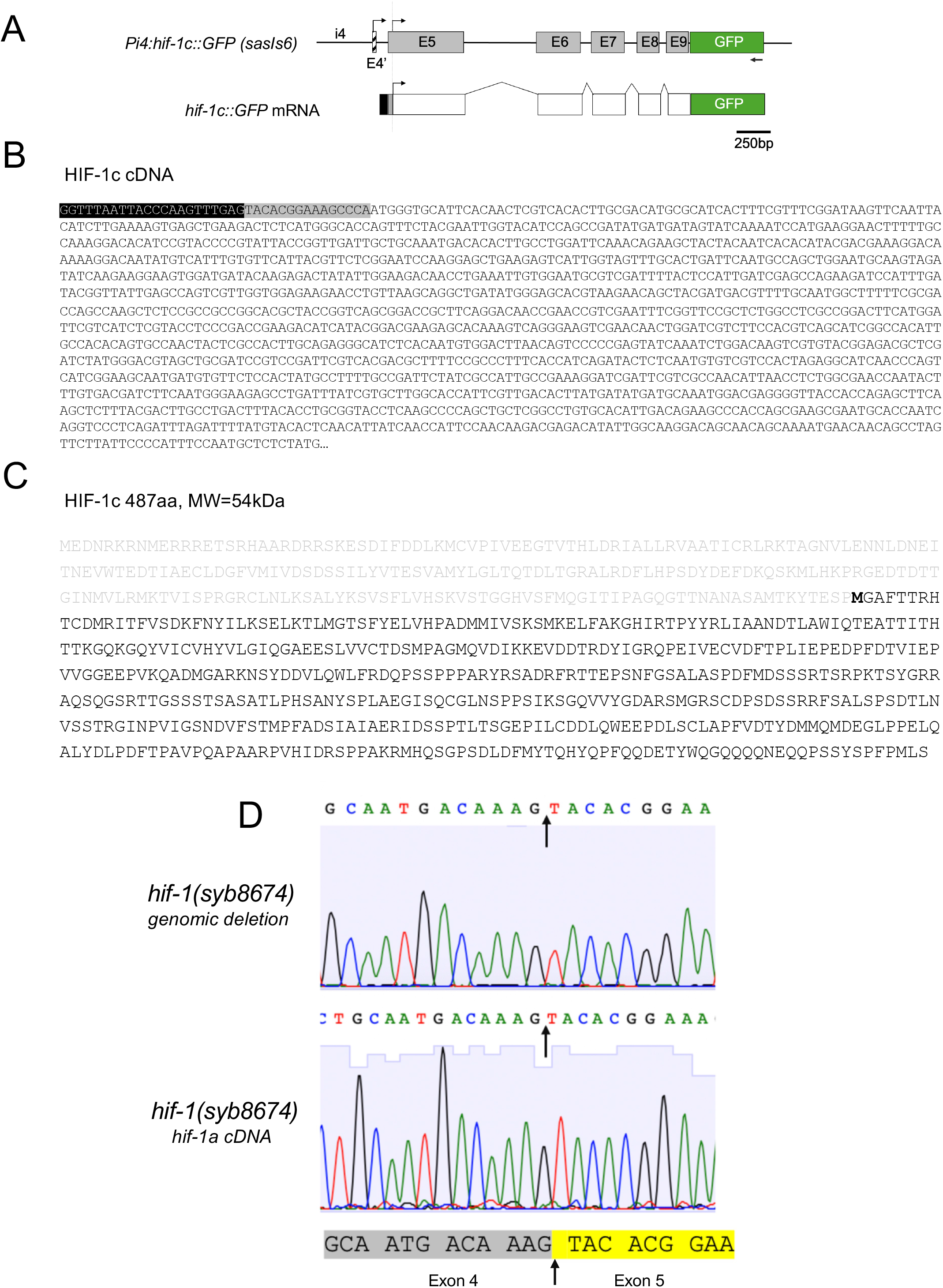
*hif-1c c*DNA analysis. (A) Schematic of the *Pi4::hif-1c::GFP* transgene. Putative start codons in frame with the *hif-1* ORF are shown with black arrows above. Diagram of cDNA from worms expressing *Pi4::hif-1c::GFP*. A 5’ SL1 primer and a GFP-specific 3’UTR primer (arrow at bottom) were used for amplification. The black box represents the SL1 linker trans-spliced into the 5’ end of the transcript highlighted in (B). (B) Sequencing of *hif-1c::GFP* cDNA revealed a SL1 sequence trans-spliced downstream of the predicted E1c (black highlight), suggesting that the native *hif-1c* mRNA likely does not use E1c for translation. A 5’UTR composed of the first untranslated nucleotides of exon 5 (grey highlight) precedes the first ATG codon used by *hif-1c*. (C) Conceptual translation of *hif-1c* mRNA. The *hif-1c* start codon, Met233, is shown in bold; grey residues indicate the N-terminal residues of HIF-1a that are not present in HIF-1c. (D) Chromatogram of *hif-1* exon 4 and 5 boundaries in *hif-1(syb8674)* genomic DNA showing the deletion breakpoint (top, arrows) and the *hif-1* cDNA produced from *syb8674* worms. The boundaries between exons 4 and 5 are highlighted in grey and yellow, respectively.

**FIGURE S5:**
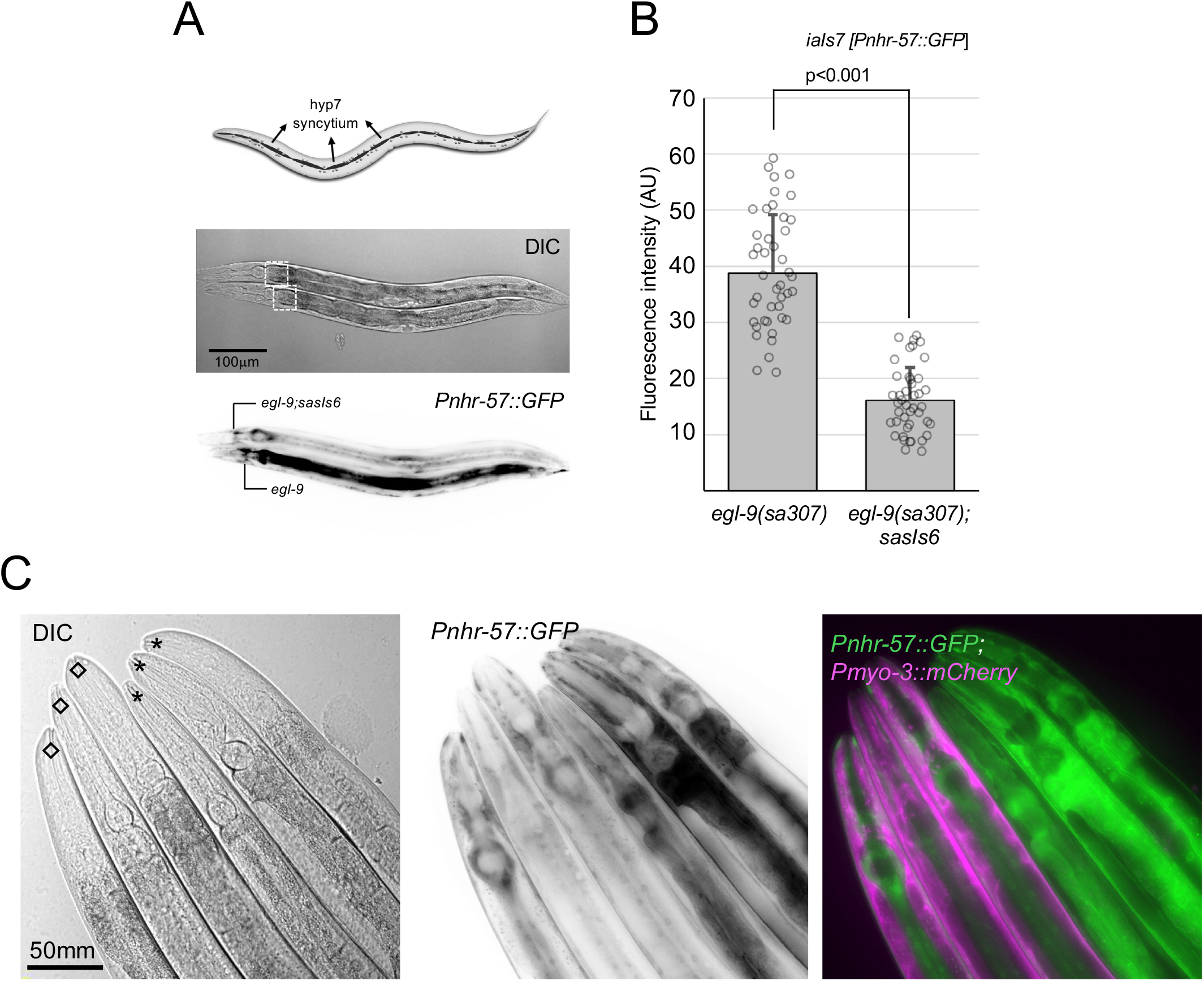
HIF-1c downregulates *Pnhr-57::GFP* expression in the hypodermis of *egl-9* mutants. (A) Representative images of *egl-9(sa307*) (bottom) and *egl-9(sa307); sasIs6* worms expressing the HIF-1 reporter *Pnhr-57::GFP (iaIs7)*. A diagram of the hypodermis is shown on top (modified from Wormbook®). Boxes in the DIC images point to the hyp7 syncytium region used for fluorescence measurements. (B) Quantification of the decrease in *Pnhr-57::GFP* expression in *egl-9(sa307); sasIs6* worms. (C) Representative images of worms expressing *Pnhr-57::GFP.* Asterisks mark the three worms expressing *Pnhr-57::GFP* alone, and diamonds point to the three worms overexpressing HIF-1c as well (*sasIs6*; marked by *Pmyo-3::mCherry*) with reduced GFP signal in the hypodermis.

**FIGURE S6:**
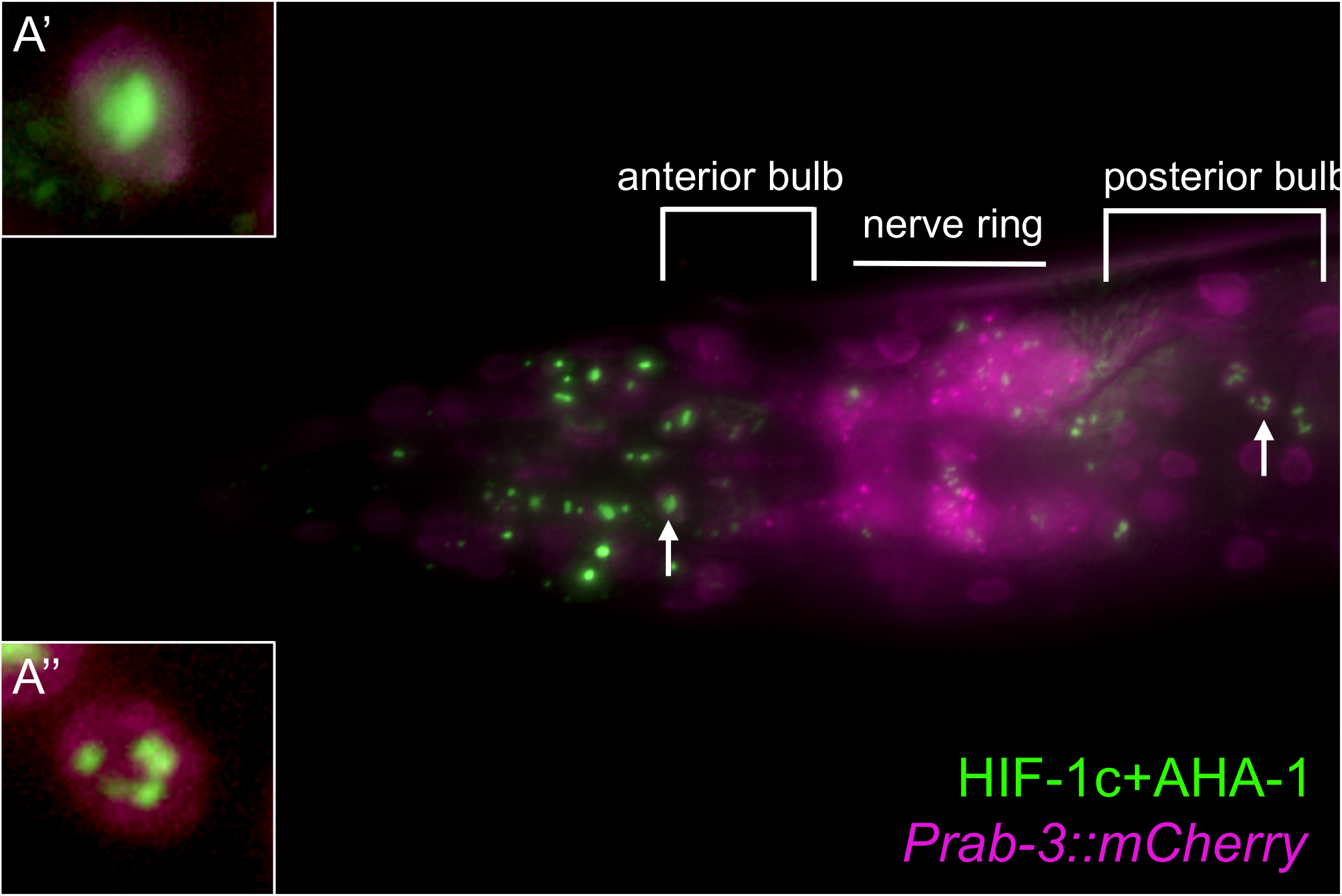
HIF-1c/AHA-1 interaction. (A) Head section of a worm expressing HIF-1C+AHA-1 BiFC constructs. Reconstituted Venus signal is shown in green. Neurons are labeled with *Prab-3::mCherry.* Insets A’ and A’’ show HIF-1c+AHA-1 aggregates in the nuclei of neurons in the anterior ganglion and retrovesicular ganglion, respectively (arrows).

**FIGURE S7:**
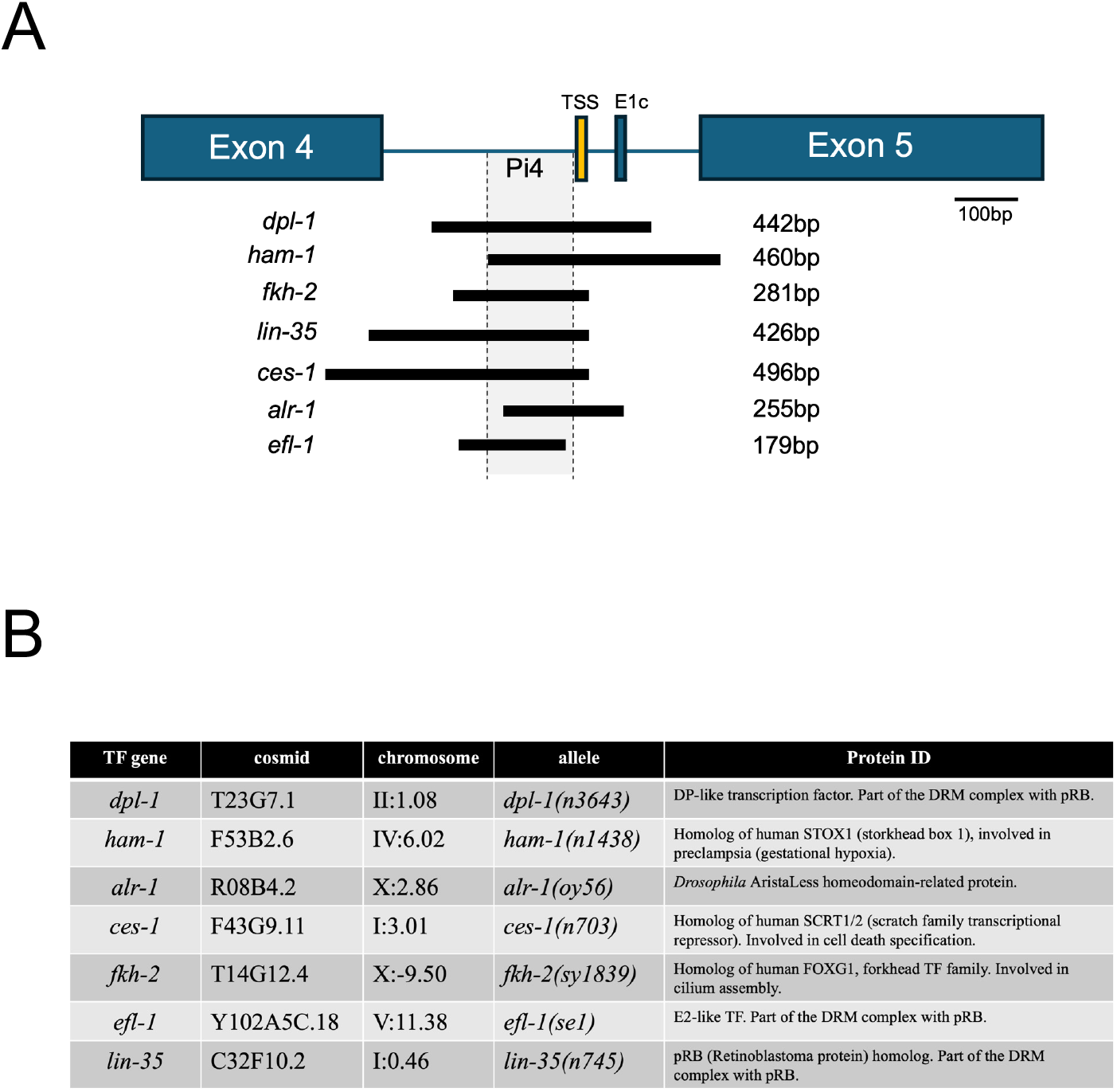
Candidate *hif-1c* transcriptional regulators. (A) Schematic of the *Pi4* region within *hif-1* intron 4. The region spanned by the primary pulldown ChIP-seq reads (WormBase® genome browser) are shown as bars underneath. Abbreviations: TSS – transcription start site; E1c – cryptic exon 1 of *hif-1c*. (B) Table of the candidate genes mentioned above, corresponding mutants and predicted protein functions.

**FIGURE S8:**
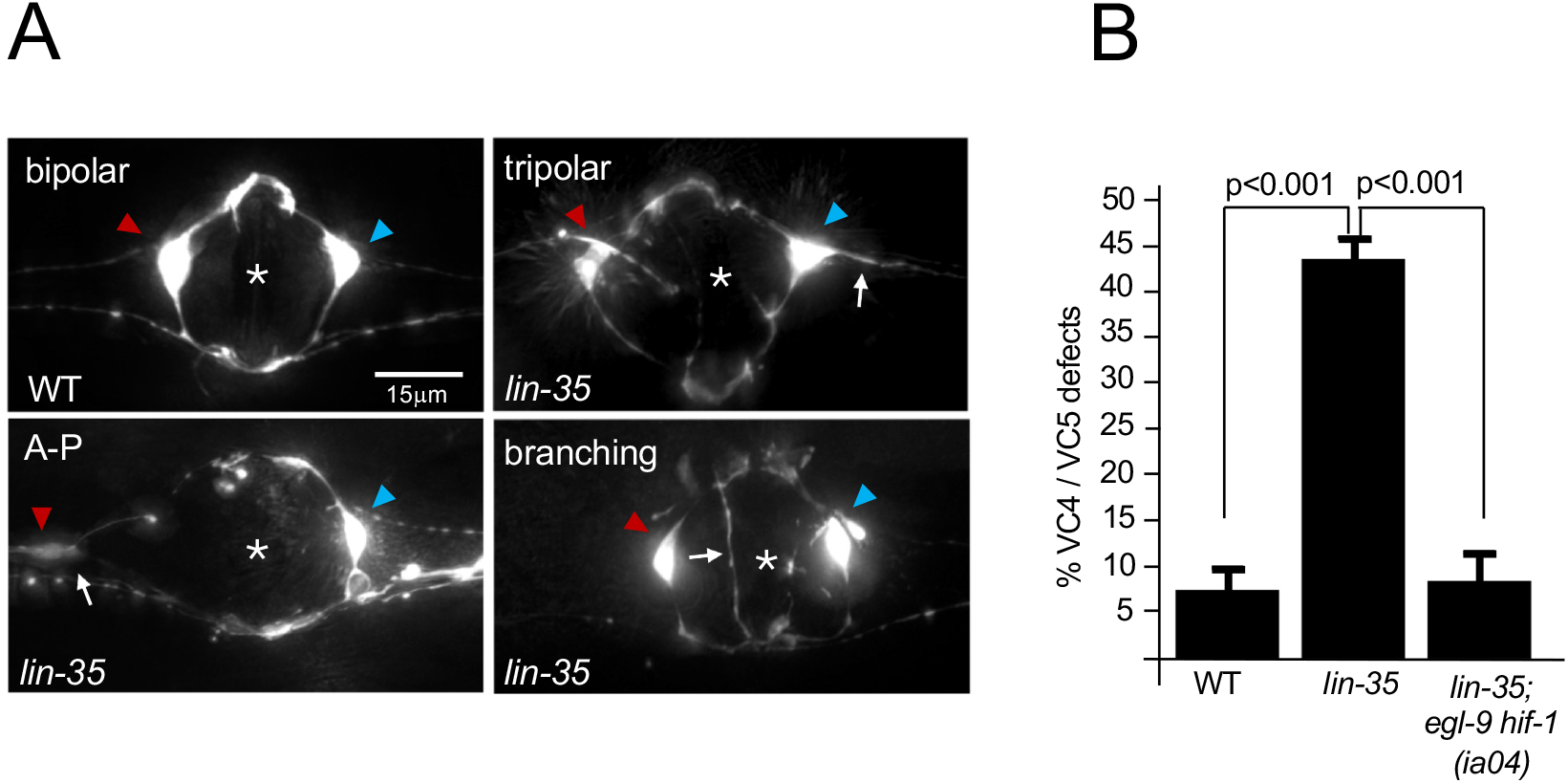
*lin-35(n745)* mutants have VC4/VC5 connectivity defects. (A) Representative images showing VC4/VC5 defects (arrows) in *lin-35* mutants, compared with the wild-type bipolar configuration of these neurons. VC4/VC5 are labeled with the *Pcat-1::GFP* reporter. VC4 is indicated by a red arrowhead, and VC5 by a blue arrowhead. Quantification of the defect is shown in B; n = 180 for each genotype.

**FIGURE S9:**
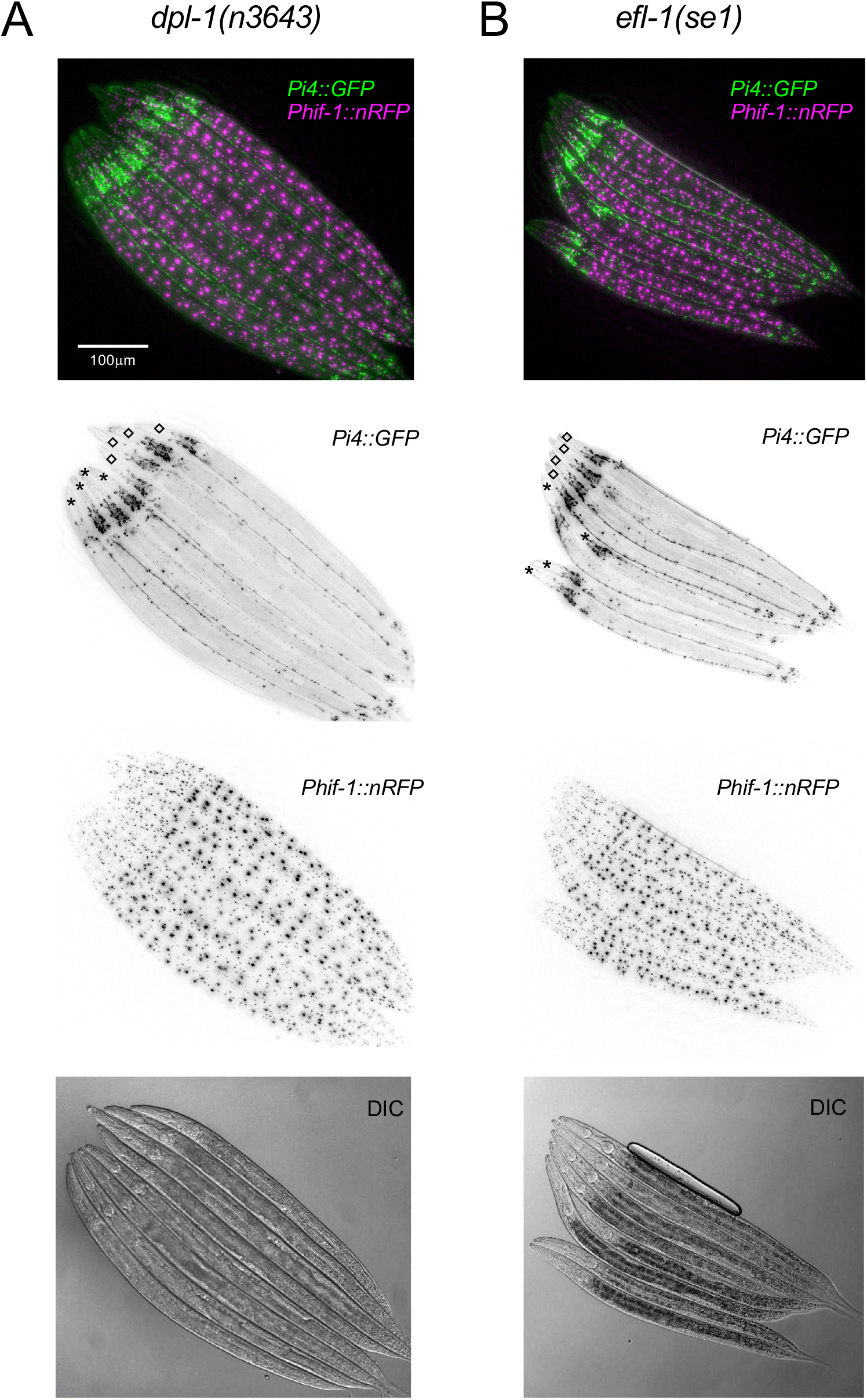
*dpl-1(n3643) and efl-1(se1)* mutants show overtly normal expression levels of *Phif-1::nRFP* and *Pi4::GFP.* Diamonds indicate control *sasIs7* worms; asterisks indicate *dpl-1* (B) and *efl-1* (B) mutants expressing *sasIs7*.

**Video S1:** Expression of HIF-1c in head neurons. Z-stack through the head of a L4 *egl-9(sa307)* animal expressing HIF-1c::GFP. GFP signal is visible in the cell bodies of these cells.

**Video S2:** Changes in HIF-1c::GFP levels in *egl-9* worms. Video of an *egl-9(sa307)* worm (background) with the overlaid anterior ganglia region (box) of an *egl-9(+)* L4 worm expressing HIF-1c::GFP. The movie starts with the *egl-9(sa307)* profile, then switches to HIF-1c::GFP expression in *egl-9(+)* worms, with neurons in the same anatomical position. The process is repeated once in the video.

**Video S3: 3D** rotation of a DAPI-stained (magenta) nucleus of an intestinal cell from a worm expressing HIF-1a + AHA-1 BiFC constructs (green).

**Video S4: 3D** rotation of a DAPI-stained (magenta) nucleus of an intestinal cell from a worm expressing HIF-1c + AHA-1 BiFC constructs (green).

## Notes

### Competing Interest Statement

The authors have declared no competing interest.

## References

1. Erecinska, M., and Silver, I.A. (2001). Tissue oxygen tension and brain sensitivity to hypoxia. Respir Physiol 128, 263–276. 10.1016/s0034-5687(01)00306-1.

2. Harris, J.J., Jolivet, R., and Attwell, D. (2012). Synaptic energy use and supply. Neuron 75, 762–777. 10.1016/j.neuron.2012.08.019.

3. Schofield, C.J., and Ratcliffe, P.J. (2004). Oxygen sensing by HIF hydroxylases. Nat Rev Mol Cell Biol 5, 343–354. 10.1038/nrm1366.

4. Semenza, G.L. (2001). HIF-1 and mechanisms of hypoxia sensing. Curr Opin Cell Biol 13, 167–171. 10.1016/s0955-0674(00)00194-0.

5. Depping, R., Hagele, S., Wagner, K.F., Wiesner, R.J., Camenisch, G., Wenger, R.H., and Katschinski, D.M. (2004). A dominant-negative isoform of hypoxia-inducible factor-1 alpha specifically expressed in human testis. Biol Reprod 71, 331–339. 10.1095/biolreprod.104.027797.

6. Maynard, M.A., Evans, A.J., Hosomi, T., Hara, S., Jewett, M.A., and Ohh, M. (2005). Human HIF-3alpha4 is a dominant-negative regulator of HIF-1 and is down-regulated in renal cell carcinoma. FASEB J 19, 1396–1406. 10.1096/fj.05-3788com.

7. Makino, Y., Cao, R., Svensson, K., Bertilsson, G., Asman, M., Tanaka, H., Cao, Y., Berkenstam, A., and Poellinger, L. (2001). Inhibitory PAS domain protein is a negative regulator of hypoxia-inducible gene expression. Nature 414, 550–554. 10.1038/35107085.

8. Pocock, R., and Hobert, O. (2008). Oxygen levels affect axon guidance and neuronal migration in Caenorhabditis elegans. Nat Neurosci 11, 894–900. 10.1038/nn.2152.

9. Stevenson, T.J., Trinh, T., Kogelschatz, C., Fujimoto, E., Lush, M.E., Piotrowski, T., Brimley, C.J., and Bonkowsky, J.L. (2012). Hypoxia disruption of vertebrate CNS pathfinding through ephrinB2 Is rescued by magnesium. PLoS Genet 8, e1002638. 10.1371/journal.pgen.1002638.

10. Bonkowsky, J.L., and Son, J.H. (2018). Hypoxia and connectivity in the developing vertebrate nervous system. Dis Model Mech 11. 10.1242/dmm.037127.

11. Jiang, H., Guo, R., and Powell-Coffman, J.A. (2001). The Caenorhabditis elegans hif-1 gene encodes a bHLH-PAS protein that is required for adaptation to hypoxia. Proc Natl Acad Sci U S A 98, 7916–7921. 10.1073/pnas.141234698.

12. Shen, C., and Powell-Coffman, J.A. (2003). Genetic analysis of hypoxia signaling and response in C elegans. Ann N Y Acad Sci 995, 191–199. 10.1111/j.1749-6632.2003.tb03222.x.

13. Saito, T.L., Hashimoto, S., Gu, S.G., Morton, J.J., Stadler, M., Blumenthal, T., Fire, A., and Morishita, S. (2013). The transcription start site landscape of C. elegans. Genome Res 23, 1348–1361. 10.1101/gr.151571.112.

14. Lee, J.W., Bae, S.H., Jeong, J.W., Kim, S.H., and Kim, K.W. (2004). Hypoxia-inducible factor (HIF-1)alpha: its protein stability and biological functions. Exp Mol Med 36, 1-12. 10.1038/emm.2004.1.

15. Maxeiner, S., Grolleman, J., Schmid, T., Kammenga, J., and Hajnal, A. (2019). The hypoxia-response pathway modulates RAS/MAPK-mediated cell fate decisions in Caenorhabditis elegans. Life Sci Alliance 2. 10.26508/lsa.201800255.

16. Ma, D.K., Vozdek, R., Bhatla, N., and Horvitz, H.R. (2012). CYSL-1 interacts with the O2-sensing hydroxylase EGL-9 to promote H2S-modulated hypoxia-induced behavioral plasticity in C. elegans. Neuron 73, 925–940. 10.1016/j.neuron.2011.12.037.

17. Pocock, R., and Hobert, O. (2010). Hypoxia activates a latent circuit for processing gustatory information in C. elegans. Nat Neurosci 13, 610–614. 10.1038/nn.2537.

18. Trent, C., Tsung, N., and Horvitz, H.R. (1983). Egg-laying defective mutants of the nematode Caenorhabditis elegans. Genetics 104, 619–647. 10.1093/genetics/104.4.619.

19. Chang, A.J., and Bargmann, C.I. (2008). Hypoxia and the HIF-1 transcriptional pathway reorganize a neuronal circuit for oxygen-dependent behavior in Caenorhabditis elegans. Proc Natl Acad Sci U S A 105, 7321–7326. 10.1073/pnas.0802164105.

20. Gort, E.H., van Haaften, G., Verlaan, I., Groot, A.J., Plasterk, R.H., Shvarts, A., Suijkerbuijk, K.P., van Laar, T., van der Wall, E., Raman, V., et al. (2008). The TWIST1 oncogene is a direct target of hypoxia-inducible factor-2alpha. Oncogene 27, 1501–1510. 10.1038/sj.onc.1210795.

21. Adler, C.E., Fetter, R.D., and Bargmann, C.I. (2006). UNC-6/Netrin induces neuronal asymmetry and defines the site of axon formation. Nat Neurosci 9, 511–518. 10.1038/nn1666.

22. Colavita, A., and Tessier-Lavigne, M. (2003). A Neurexin-related protein, BAM-2, terminates axonal branches in C. elegans. Science 302, 293–296. 10.1126/science.1089163.

23. Habibi-Babadi, N., Su, A., de Carvalho, C.E., and Colavita, A. (2010). The N-glycanase png-1 acts to limit axon branching during organ formation in Caenorhabditis elegans. J Neurosci 30, 1766–1776. 10.1523/JNEUROSCI.4962-08.2010.

24. Sanchez-Alvarez, L., Visanuvimol, J., McEwan, A., Su, A., Imai, J.H., and Colavita, A. (2011). VANG-1 and PRKL-1 cooperate to negatively regulate neurite formation in Caenorhabditis elegans. PLoS Genet 7, e1002257. 10.1371/journal.pgen.1002257.

25. Boulin, T., Pocock, R., and Hobert, O. (2006). A novel Eph receptor-interacting IgSF protein provides C. elegans motoneurons with midline guidepost function. Curr Biol 16, 1871–1883. 10.1016/j.cub.2006.08.056.

26. Heikkila, M., Pasanen, A., Kivirikko, K.I., and Myllyharju, J. (2011). Roles of the human hypoxia-inducible factor (HIF)-3alpha variants in the hypoxia response. Cell Mol Life Sci 68, 3885–3901. 10.1007/s00018-011-0679-5.

27. Niu, W., Lu, Z.J., Zhong, M., Sarov, M., Murray, J.I., Brdlik, C.M., Janette, J., Chen, C., Alves, P., Preston, E., et al. (2011). Diverse transcription factor binding features revealed by genome-wide ChIP-seq in C. elegans. Genome Res 21, 245–254. 10.1101/gr.114587.110.

28. Wang, D., Kennedy, S., Conte, D., Jr., Kim, J.K., Gabel, H.W., Kamath, R.S., Mello, C.C., and Ruvkun, G. (2005). Somatic misexpression of germline P granules and enhanced RNA interference in retinoblastoma pathway mutants. Nature 436, 593–597. 10.1038/nature04010.

29. Tops, B.B., Tabara, H., Sijen, T., Simmer, F., Mello, C.C., Plasterk, R.H., and Ketting, R.F. (2005). RDE-2 interacts with MUT-7 to mediate RNA interference in Caenorhabditis elegans. Nucleic Acids Res 33, 347–355. 10.1093/nar/gki183.

30. Ceol, C.J., and Horvitz, H.R. (2001). dpl-1 DP and efl-1 E2F act with lin-35 Rb to antagonize Ras signaling in C. elegans vulval development. Mol Cell 7, 461–473. 10.1016/s1097-2765(01)00194-0.

31. Ravenna, L., Salvatori, L., and Russo, M.A. (2016). HIF3alpha: the little we know. FEBS J 283, 993–1003. 10.1111/febs.13572.

32. Yamashita, T., Ohneda, O., Nagano, M., Iemitsu, M., Makino, Y., Tanaka, H., Miyauchi, T., Goto, K., Ohneda, K., Fujii-Kuriyama, Y., et al. (2008). Abnormal heart development and lung remodeling in mice lacking the hypoxia-inducible factor-related basic helix-loop-helix PAS protein NEPAS. Mol Cell Biol 28, 1285–1297. 10.1128/MCB.01332-07.

33. Lee, H.C., and Tsai, S.J. (2017). Endocrine targets of hypoxia-inducible factors. J Endocrinol 234, R53–R65. 10.1530/JOE-16-0653.

34. Wu, Z., Ghosh-Roy, A., Yanik, M.F., Zhang, J.Z., Jin, Y., and Chisholm, A.D. (2007). Caenorhabditis elegans neuronal regeneration is influenced by life stage, ephrin signaling, and synaptic branching. Proc Natl Acad Sci U S A 104, 15132–15137. 10.1073/pnas.0707001104.

35. Kalichamy, K., Arnold, M., Hill, T., Johnson, A., McCardel, K., Taylor, W.K., Upton, B., and Hudson, M.L. (2025). The vab-1/Eph receptor and ephrins efn-1 and efn-4 are required for C. elegans chemosensory and thermosensory circuit assembly and function. BMC Neurosci 26, 65. 10.1186/s12868-025-00983-3.

36. Mohamed, A.M., and Chin-Sang, I.D. (2006). Characterization of loss-of-function and gain-of-function Eph receptor tyrosine kinase signaling in C. elegans axon targeting and cell migration. Dev Biol 290, 164–176. 10.1016/j.ydbio.2005.11.019.

37. Talluri, S., and Dick, F.A. (2012). Regulation of transcription and chromatin structure by pRB: here, there and everywhere. Cell Cycle 11, 3189–3198. 10.4161/cc.21263.

38. Lee, J.S., Thomas, D.M., Gutierrez, G., Carty, S.A., Yanagawa, S., and Hinds, P.W. (2006). HES1 cooperates with pRb to activate RUNX2-dependent transcription. J Bone Miner Res 21, 921–933. 10.1359/jbmr.060303.

39. Kirienko, N.V., and Fay, D.S. (2007). Transcriptome profiling of the C. elegans Rb ortholog reveals diverse developmental roles. Dev Biol 305, 674–684. 10.1016/j.ydbio.2007.02.021.

40. Amellem, O., Sandvik, J.A., Stokke, T., and Pettersen, E.O. (1998). The retinoblastoma protein-associated cell cycle arrest in S-phase under moderate hypoxia is disrupted in cells expressing HPV18 E7 oncoprotein. Br J Cancer 77, 862–872. 10.1038/bjc.1998.143.

41. Budde, A., Schneiderhan-Marra, N., Petersen, G., and Brune, B. (2005). Retinoblastoma susceptibility gene product pRB activates hypoxia-inducible factor-1 (HIF-1). Oncogene 24, 1802–1808. 10.1038/sj.onc.1208369.

42. Labrecque, M.P., Takhar, M.K., Jagdeo, J.M., Tam, K.J., Chiu, C., Wang, T.Y., Prefontaine, G.G., Cox, M.E., and Beischlag, T.V. (2014). A TRIP230-retinoblastoma protein complex regulates hypoxia-inducible factor-1alpha-mediated transcription and cancer cell invasion. PLoS One 9, e99214. 10.1371/journal.pone.0099214.

43. To, C., Farnsworth, R.H., Vail, M.E., Chheang, C., Gargett, C.E., Murone, C., Llerena, C., Major, A.T., Scott, A.M., Janes, P.W., and Lackmann, M. (2014). Hypoxia-controlled EphA3 marks a human endometrium-derived multipotent mesenchymal stromal cell that supports vascular growth. PLoS One 9, e112106. 10.1371/journal.pone.0112106.

44. Husain, A., Chiu, Y.T., Sze, K.M., Ho, D.W., Tsui, Y.M., Suarez, E.M.S., Zhang, V.X., Chan, L.K., Lee, E., Lee, J.M., et al. (2022). Ephrin-A3/EphA2 axis regulates cellular metabolic plasticity to enhance cancer stemness in hypoxic hepatocellular carcinoma. J Hepatol 77, 383–396. 10.1016/j.jhep.2022.02.018.

45. Sheikh, A., Meng, X., Liu, J., Mikhailova, A., Kao, J.P.Y., McQuillen, P.S., and Kanold, P.O. (2019). Neonatal Hypoxia-Ischemia Causes Functional Circuit Changes in Subplate Neurons. Cereb Cortex 29, 765–776. 10.1093/cercor/bhx358.

46. Lai, M.C., and Yang, S.N. (2011). Perinatal hypoxic-ischemic encephalopathy. J Biomed Biotechnol 2011, 609813. 10.1155/2011/609813.

## References

1. Nalivaeva, N.N., Turner, A.J., and Zhuravin, I.A. (2018). Role of Prenatal Hypoxia in Brain Development, Cognitive Functions, and Neurodegeneration. Front Neurosci 12, 825. 10.3389/fnins.2018.00825.

2. Wang, B., Zeng, H., Liu, J., and Sun, M. (2021). Effects of Prenatal Hypoxia on Nervous System Development and Related Diseases. Frontiers in Neuroscience Volume 15 - 2021. 10.3389/fnins.2021.755554.

3. Semenza, G.L., and Wang, G.L. (1992). A nuclear factor induced by hypoxia via de novo protein synthesis binds to the human erythropoietin gene enhancer at a site required for transcriptional activation. Mol Cell Biol 12, 5447–5454. 10.1128/mcb.12.12.5447-5454.1992.

4. Wang, G.L., and Semenza, G.L. (1993). Characterization of hypoxia-inducible factor 1 and regulation of DNA binding activity by hypoxia. J Biol Chem 268, 21513–21518.

5. Stevenson, T.J., Trinh, T., Kogelschatz, C., Fujimoto, E., Lush, M.E., Piotrowski, T., Brimley, C.J., and Bonkowsky, J.L. (2012). Hypoxia disruption of vertebrate CNS pathfinding through ephrinB2 Is rescued by magnesium. PLoS Genet 8, e1002638. 10.1371/journal.pgen.1002638.

6. Pocock, R., and Hobert, O. (2008). Oxygen levels affect axon guidance and neuronal migration in Caenorhabditis elegans. Nat Neurosci 11, 894–900. 10.1038/nn.2152.

7. Arias, C.F., Acosta, F.J., Bertocchini, F., and Fernández-Arias, C. (2025). Redefining the role of hypoxia-inducible factors (HIFs) in oxygen homeostasis. Commun Biol 8, 446. 10.1038/s42003-025-07896-1.

8. Ding, D., Zhang, J., Du, B., Wang, X., Hou, L., Guo, S., Chen, B., and Kang, L. (2022). Non-canonical function of an Hif-1α splice variant contributes to the sustained flight of locusts. eLife 11, e74554. 10.7554/eLife.74554.

9. Angeles-Albores, D., Puckett Robinson, C., Williams, B.A., Wold, B.J., and Sternberg, P.W. (2018). Reconstructing a metazoan genetic pathway with transcriptome-wide epistasis measurements. Proceedings of the National Academy of Sciences 115, E2930–E2939. doi:10.1073/pnas.1712387115.

10. Fujita, N., Chiba, K., Shapiro, I.M., and Risbud, M.V. (2012). HIF-1α and HIF-2α degradation is differentially regulated in nucleus pulposus cells of the intervertebral disc. Journal of Bone and Mineral Research 27, 401–412. 10.1002/jbmr.538.

11. Maxeiner, S., Grolleman, J., Schmid, T., Kammenga, J., and Hajnal, A. (2019). The hypoxia-response pathway modulates RAS/MAPK-mediated cell fate decisions in Caenorhabditis elegans. Life Sci Alliance 2. 10.26508/lsa.201800255.

12. Yoshina, S., Suehiro, Y., Kage-Nakadai, E., and Mitani, S. (2016). Locus-specific integration of extrachromosomal transgenes in C. elegans with the CRISPR/Cas9 system. Biochemistry and biophysics reports 5, 70–76.

13. Mello, C.C., Kramer, J.M., Stinchcomb, D., and Ambros, V. (1991). Efficient gene transfer in C.elegans: extrachromosomal maintenance and integration of transforming sequences. Embo j 10, 3959–3970. 10.1002/j.1460-2075.1991.tb04966.x.

14. Zhou, S., and Chen, L. (2011). Neural integrity is maintained by dystrophin in C. elegans. Journal of Cell Biology 192, 349–363. 10.1083/jcb.201006109.

15. Schindelin, J., Arganda-Carreras, I., Frise, E., Kaynig, V., Longair, M., Pietzsch, T., Preibisch, S., Rueden, C., Saalfeld, S., Schmid, B., et al. (2012). Fiji: an open-source platform for biological-image analysis. Nat Methods 9, 676–682. 10.1038/nmeth.2019.

16. Harris, T.W., Antoshechkin, I., Bieri, T., Blasiar, D., Chan, J., Chen, W.J., De La Cruz, N., Davis, P., Duesbury, M., Fang, R., et al. (2010). WormBase: a comprehensive resource for nematode research. Nucleic Acids Res 38, D463–467. 10.1093/nar/gkp952.

17. Brenner, S. (1974). The genetics of Caenorhabditis elegans. Genetics 77, 71–94. 10.1093/genetics/77.1.71.

18. Mello, C., and Fire, A. (1995). Chapter 19 DNA Transformation. In Methods in Cell Biology, H.F. Epstein, and D.C. Shakes, eds. (Academic Press), pp. 451–482. 10.1016/S0091-679X(08)61399-0.

19. Mariol, M.C., Walter, L., Bellemin, S., and Gieseler, K. (2013). A rapid protocol for integrating extrachromosomal arrays with high transmission rate into the C. elegans genome. J Vis Exp, e50773. 10.3791/50773.

20. Hall, D.H., and Russell, R.L. (1991). The posterior nervous system of the nematode Caenorhabditis elegans: serial reconstruction of identified neurons and complete pattern of synaptic interactions. J Neurosci 11, 1–22. 10.1523/jneurosci.11-01-00001.1991.

21. Pocock, R., and Hobert, O. (2010). Hypoxia activates a latent circuit for processing gustatory information in C. elegans. Nat Neurosci 13, 610–614. 10.1038/nn.2537.

22. Hiatt, S.M., Shyu, Y.J., Duren, H.M., and Hu, C.D. (2008). Bimolecular fluorescence complementation (BiFC) analysis of protein interactions in Caenorhabditis elegans. Methods 45, 185–191. 10.1016/j.ymeth.2008.06.003.

23. Shyu, Y.J., Hiatt, S.M., Duren, H.M., Ellis, R.E., Kerppola, T.K., and Hu, C.D. (2008). Visualization of protein interactions in living Caenorhabditis elegans using bimolecular fluorescence complementation analysis. Nat Protoc 3, 588–596. 10.1038/nprot.2008.16.

24. Glenwinkel, L., Taylor, S.R., Langebeck-Jensen, K., Pereira, L., Reilly, M.B., Basavaraju, M., Rafi, I., Yemini, E., Pocock, R., Sestan, N., et al. (2021). In silico analysis of the transcriptional regulatory logic of neuronal identity specification throughout the C. elegans nervous system. Elife 10. 10.7554/eLife.64906.

25. Brdlik, C.M., Niu, W., and Snyder, M. (2014). Chromatin immunoprecipitation and multiplex sequencing (ChIP-Seq) to identify global transcription factor binding sites in the nematode Caenorhabditis elegans. Methods Enzymol 539, 89–111. 10.1016/b978-0-12-420120-0.00007-4.

26. Niu, W., Lu, Z.J., Zhong, M., Sarov, M., Murray, J.I., Brdlik, C.M., Janette, J., Chen, C., Alves, P., Preston, E., et al. (2011). Diverse transcription factor binding features revealed by genome-wide ChIP-seq in C. elegans. Genome Res 21, 245–254. 10.1101/gr.114587.110.

27. Ren, B., Robert, F., Wyrick, J.J., Aparicio, O., Jennings, E.G., Simon, I., Zeitlinger, J., Schreiber, J., Hannett, N., Kanin, E., et al. (2000). Genome-wide location and function of DNA binding proteins. Science 290, 2306–2309. 10.1126/science.290.5500.2306.

